# WRN helicase upregulates mitophagy by resolving an intricate nexus of G-quadruplexes-R loops-ATG7 pre-mRNA maturation in cancer

**DOI:** 10.64898/2025.12.11.692796

**Authors:** Pooja Gupta, Mrityunjay Tyagi, Ananda Guha Majumdar, Ganesh Pai Bellare, Saikat Chakraborty, Shambhavi Mishra, Megha Tawate, Avik Chakraborty, Sujit Kumar Bhutia, Birija Sankar Patro

## Abstract

Werner syndrome (WS) is a progeroid and cancer-predisposition disorder caused by loss of the Werner RECQ helicase-exonuclease (WRN), a key genome-maintenance enzyme essential for replication-stress signalling and DNA-repair. WS patients also develop metabolic abnormalities, including fatty-liver and diabetes, suggesting a link between WRN-deficiency and mitochondrial-dysfunction. WRN is frequently epigenetically silenced in cancers, yet its precise role in mitochondrial homeostasis in cancer remains unclear. Here, we define a role for WRN in regulating mitophagy and autophagy in cancer. WRN-deficient cells show defective mitochondrial respiration, morphology, and mitophagosome/autophagosome-maturation under basal and cisplatin-induced stress. Mechanistically, WRN-loss causes strong reduction of ATG7-protein, compromising autophagosome-biogenesis. Chromatin immunoprecipitation reveals accumulation of unresolved G-quadruplex structures (G4-DNA) across the ATG7-locus in WRN-deficient cells. Paradoxically, ATG7-mRNA expression is elevated despite reduced ATG7-protein in WRN-deficient cells, indicating a post-transcriptional defect. Further, we show that WRN resolves G4-DNA which prevent R-loops formation and interacts with the mRNA-processing factor U2AF35, independent of its helicase-exonuclease functions, to promote maturation of nascent ATG7 transcripts. In cancer patients, WRN level also inversely correlated with post-transcriptional defects in ATG7 mRNA. Collectively, our findings suggest pivotal association of WRN-loss in autophagy fidelity, which may further contribute to oncogenic transformation in WRN-deficient tissues and exacerbate cancer susceptibility in WS-patients.

**Significance statement:** WRN is well recognized for its roles in DNA repair and genome maintenance, which are essential for suppressing tumorigenesis and Werner syndrome (WS)–associated premature-aging. However, its functions in mitochondrial regulation remain underexplored, despite WS patients exhibiting severe metabolic defects and increased cancer risk. Here, we uncover a mechanistic link between WRN and autophagy/mitophagy, showing that WRN resolves G-quadruplexes and R-loops to enable proper post-transcriptional processing and translation of ATG7, a key autophagy enzyme. WRN-loss associated defective-autophagy may heighten the initiation of tumorigenesis in both WS-patients and in normal individual with mutated WRN in different tissue-types. As WRN is actively pursued as a synthetic-lethal target and multiple WRN inhibitors progress through clinical-development, our findings highlight mitochondrial quality-control defects as an additional determinant of WRN-targeted therapeutic-response.

## Introduction

Werner syndrome (WS), a prototypical premature aging disorder, results from loss-of-function mutations in the WRN gene, which maps to chromosome 8p12 ^1^. Patients with WS exhibit a markedly elevated predisposition to multiple malignancies, including thyroid carcinoma, melanoma, breast cancer, meningioma, and soft tissue and bone sarcomas ^1^. In line with these clinical observations, WRN deficiency or reduced WRN expression has been documented across diverse human cancers, frequently arising from epigenetic silencing *via* promoter hypermethylation and, in certain cases, somatic mutations within the WRN locus ^2–4^. Loss or attenuation of WRN function bears significant prognostic implications, as diminished WRN expression correlates with aggressive molecular subtypes and poorer overall survival ^5^. Notably, tumors exhibiting concurrent WRN downregulation and topoisomerase I overexpression are associated with particularly adverse clinical outcomes ^6^.

WRN, a member of the RECQ helicase family, plays a pivotal role in preserving genomic stability. It is unique among RECQ helicases in possessing dual catalytic activities - a 3′→5′ DNA helicase and a 3′→5′ exonuclease ^7^. WRN unwinds diverse DNA secondary structures and orchestrates essential DNA metabolic processes, including replication, repair, recombination, and telomere maintenance ^8^. Through interactions with key genome stability factors such as RPA, RAD51, Ku70/80, and p53, WRN coordinates DNA damage signaling and repair responses (DDR) ^9^. Recent studies further implicate WRN in replication fork protection in BRCA2-deficient cells, in suppressing R-loop–associated DNA damage, and in stabilizing microsatellite sequences ^10^. Together with RPA, WRN facilitates replication fork recovery following replication stress, counteracting G-quadruplex (G4)–mediated fork stalling ^11^. Our previous work demonstrated that WRN orchestrates the repair of topoisomerase I-DNA covalent complexes, promoting robust single-stranded DNA generation and CHK1-mediated NF-κB activation - a process independent of both its helicase and exonuclease activities ^12,13^. Functionally, WRN loss induces replication stress, chromosomal instability, telomere attrition, and accumulation of DNA damage, culminating in premature senescence and tumorigenesis. Given its central role in genome maintenance, WRN has emerged as a synthetic lethal vulnerability, particularly in microsatellite instability (MSI)-high cancers, thereby attracting significant therapeutic interest ^14–16^. These advances on the role of WRN in nuclear DDR have driven a paradigm shift toward the development of WRN-targeting agents, several of which are currently under preclinical and clinical evaluation^17^.

Clinically, WS manifests as accelerated aging, characterized by early-onset malignancies, short stature, alopecia, premature graying, progeroid facial features, juvenile cataracts, dyslipidemia, atherosclerosis, and insulin-resistant diabetes ^1^. These metabolic abnormalities underscore the multifaceted, yet incompletely understood, physiological roles of WRN beyond genome surveillance. Recent studies have revealed that mitochondrial dysfunction and defective mitophagy contribute significantly to the accelerated aging phenotype in WS ^18^. Restoration of cellular NAD levels ameliorates these defects, improving mitochondrial homeostasis and extending lifespan in *Caenorhabditis elegans* and *Drosophila melanogaster* models of WS, largely by preserving stem cell function and delaying organismal aging ^18^. *WRN* mutant mice exhibit premature hepatic sinusoidal endothelial defenestration accompanied by inflammatory responses and metabolic dysfunction ^19^. Notably, dietary vitamin C supplementation ameliorated the reduced mean lifespan of WRN-deficient mice and reversed multiple age-associated phenotypes, including adipose tissue degeneration, hepatic endothelial defenestration, genomic instability, and systemic inflammation ^19^. At the cellular level, WRN deficiency induces endoplasmic reticulum (ER) stress and impaired autophagy in fibroblasts ^20^. Acidic domain of WRN protein is critical for autophagy and up-regulates age associated proteins ^21^. Downregulation of WRN drives a metabolic shift, disrupts mitochondrial function, and perturbs redox homeostasis, thereby restricting cancer cell proliferation ^22,23^. Although aging and cancer share overlapping molecular hallmarks, their regulatory networks diverge fundamentally. In this context, the contribution of WRN deficiency to cancer-associated adaptation for mitochondrial homeostasis remains largely unexplored. Moreover, the specific mechanistic roles of WRN, a nuclear protein, in mitophagy and autophagy regulation within cancer cells are still enigmatic and poorly defined. In the current report, we uncover an unexpected role of WRN in resolving a complex nexus of G-quadruplex DNA (G4) structures, R-loops, and pre-mRNA maturation, which is essential for the efficient expression of ATG7 and, consequently, for effective mitophagy and autophagy in cancers.

## Results

### WRN plays a pivotal role in preserving mitochondrial function and morphology

Given that Werner syndrome (WS) patients display profound metabolic abnormalities, including hepatic steatosis, diabetes, and cancer predisposition ^1^, we investigated whether WRN deficiency compromises mitochondrial function in cancer cells. To address this, WRN knockout (WRN^-/-^) U2-OS (osteosarcoma) cell line was generated using the CRISPR-Cas9 double-nickase system (Fig. 1A). Assessment of mitochondrial respiration *via* extracellular flux analysis revealed markedly reduced basal and maximal oxygen consumption rates (OCR) and ATP-linked respiration in WRN^-/-^ cells compared to WRN^+/+^ cells (Fig. 1B,C), indicating severe mitochondrial dysfunction upon WRN loss. Interestingly, glucose uptake was approximately two-fold higher in WRN^-/-^ cells relative to WRN^+/+^ cells (Fig. 1D,E), suggesting a compensatory shift toward glycolytic or alternative glucose-dependent metabolic pathways. Because PARP1-mediated PARylation (poly-ADP ribosylation) consumes NAD^+^ in response to oxidative or genotoxic stress ^24^, excessive activation of PARP1 can deplete NAD^+^ pools, thereby perturbing mitochondrial metabolism. Consistent with this, WRN^-/-^ cells exhibited elevated basal PARylation and reduced total ATP content, further supporting the occurrence of mitochondrial dysfunction in WRN-deficient cells (Fig. S1A,B). Despite these metabolic alterations, mitochondrial membrane potential, assessed using JC-1 staining, remained comparable between WRN^-/-^ and WRN^+/+^ cells (Fig. 1F). Intriguingly, mitochondrial mass, quantified by MitoTracker Green (MTG) staining, which labels mitochondria independent of membrane potential, was slightly but significantly increased in WRN^-/-^ cells (Fig. 1G,H). Morphometric analysis further demonstrated reduced mitochondrial branching and filament length in WRN^-/-^ cells relative to controls (Fig. 1I-K, Fig. S1C), indicative of fragmented mitochondrial networks. Collectively, these findings establish that WRN is essential for sustaining mitochondrial bioenergetic capacity and preserving mitochondrial morphology, and implicating a role of WRN in the regulation of basal mitochondrial homeostasis.

**Figure 1:**
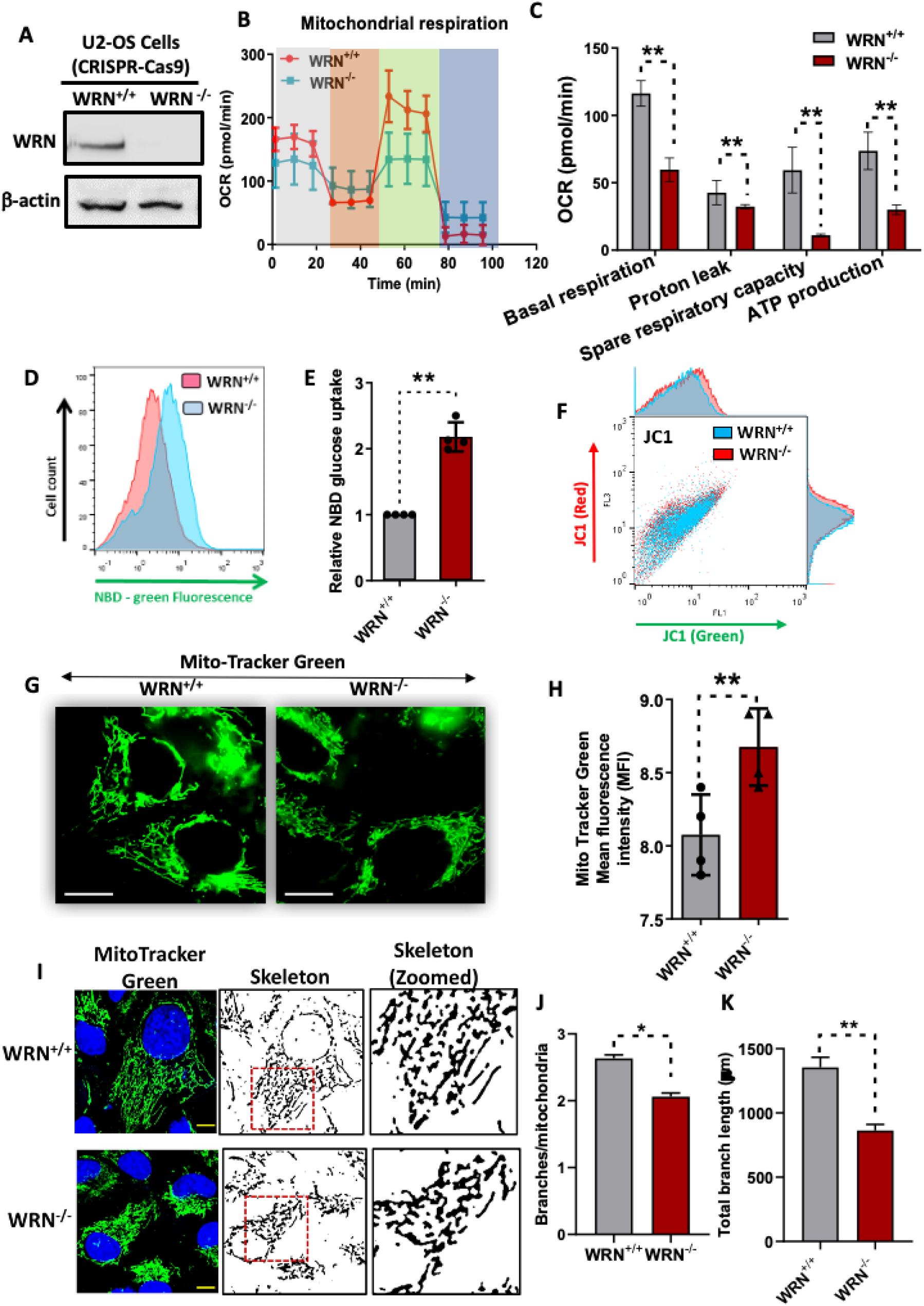
Mitochondrial function and morphology in WRN-deficient cells. (A) WRN depleted cells (WRN^-/-^) were generated by CRISPR-Cas9 double nickase plasmid system and then the expression was assessed by Western blotting. The blot was probed for WRN and β-actin as a loading control. (B, C) Mitochondrial respiration was measured in WRN^+/+^ and WRN^-/-^ cells using the Seahorse XF Analyzer. The oxygen consumption rate (OCR) was monitored over time, and phases of basal respiration, proton leak, maximal respiratory capacity and reserve capacity were indicated by grey, orange, green and blue shades, respectively in B. (D, E) NBD-glucose uptake was analyzed by flow cytometry in WRN^+/+^ and WRN^-/-^ cells, measuring the green fluorescence of NBD-glucose uptake. Relative NBD-glucose uptake was quantified to assess glucose transport in WRN^+/+^ and WRN^-/-^ cells. (F) Mitochondrial membrane potential was evaluated by flow cytometry using JC-1 dye. The fluorescence emission at red and green channels was analyzed to assess membrane potential in WRN^+/+^ and WRN^-/-^ cells. (G, H) Mitochondrial morphology was visualized using Mito-Tracker Green staining and quantified the intensity in WRN^+/+^ and WRN^-/-^ cells. Representative images of mitochondrial fluorescence are shown. (I-K) Mitochondrial morphology and branching were assessed using Mito-Tracker Green staining followed by skeletonizing of mitochondrial structures in WRN^+/+^ and WRN^-/-^ cells. Representative images are shown. Mitochondrial branching per mitochondrion and total mitochondrial branch length was quantified in WRN^+/+^ and WRN^-/-^ cells, measuring extent of mitochondrial network. All the values indicated are mean ± SEM (n = 4) \*\**p* < 0.01, \**p* <0.05.

### WRN promotes autophagosome and mitophagosome maturation

Given the essential role of WRN in maintaining basal mitochondrial function and morphology, we next investigated how WRN modulates mitochondrial homeostasis in response to mitochondrial stress induced by chemotherapeutic agents. As cisplatin (*cis*-diamminedichloroplatinum: CDDP) is widely used in clinical oncology and known to trigger mitochondrial damage through reactive oxygen species (ROS) generation, it was chosen as a model system for subsequent experiments ^25,26^. Consistent with previous reports ^25^, cisplatin treatment elicited robust cellular ROS accumulation at 24 h, which subsided by 48 h in WRN^+/+^ cells (Fig. S2A,B). In contrast, WRN^-/-^ cells exhibited persistently elevated ROS levels under identical treatment (Fig. S2A,B). Interestingly, basal and cisplatin-induced mitochondrial ROS was significantly higher in WRN-deficient cells than WRN-proficient cells (10 µM; Fig. S2C). Moreover, cisplatin treatment caused a greater accumulation of mitochondria in WRN-deficient cells, with mitochondrial mass elevated by approximately 32% in WRN^-/-^ *versus* WRN^+/+^ cells after 48 h (Fig. S2D,E). Morphometric analyses revealed that cisplatin markedly reduced mitochondrial branching and total filament length in WRN^+/+^ cells but not in WRN^-/-^ cells (Fig. 2A-C), implying attenuated mitochondrial fission dynamics in the absence of WRN. Notably, mitochondrial membrane potential (Δψm) loss, determined by JC-1 staining, was significantly higher in WRN^-/-^ cells following cisplatin exposure (Fig. 2D,E). These findings indicate that, despite increased mitochondrial mass, WRN^-/-^ cells accumulate dysfunctional mitochondria, suggestive of impaired mitophagic clearance. To validate this hypothesis, cells were co-treated with bafilomycin A1 (BafA1), an inhibitor of autophagosome-lysosome fusion, and cisplatin. BafA1 significantly enhanced mitochondrial accumulation in WRN^+/+^ cells relative to cisplatin alone, whereas no comparable effect was observed in WRN^-/-^ cells (Fig. S2F), confirming a functional role of WRN in mitophagy progression.

**Figure 2:**
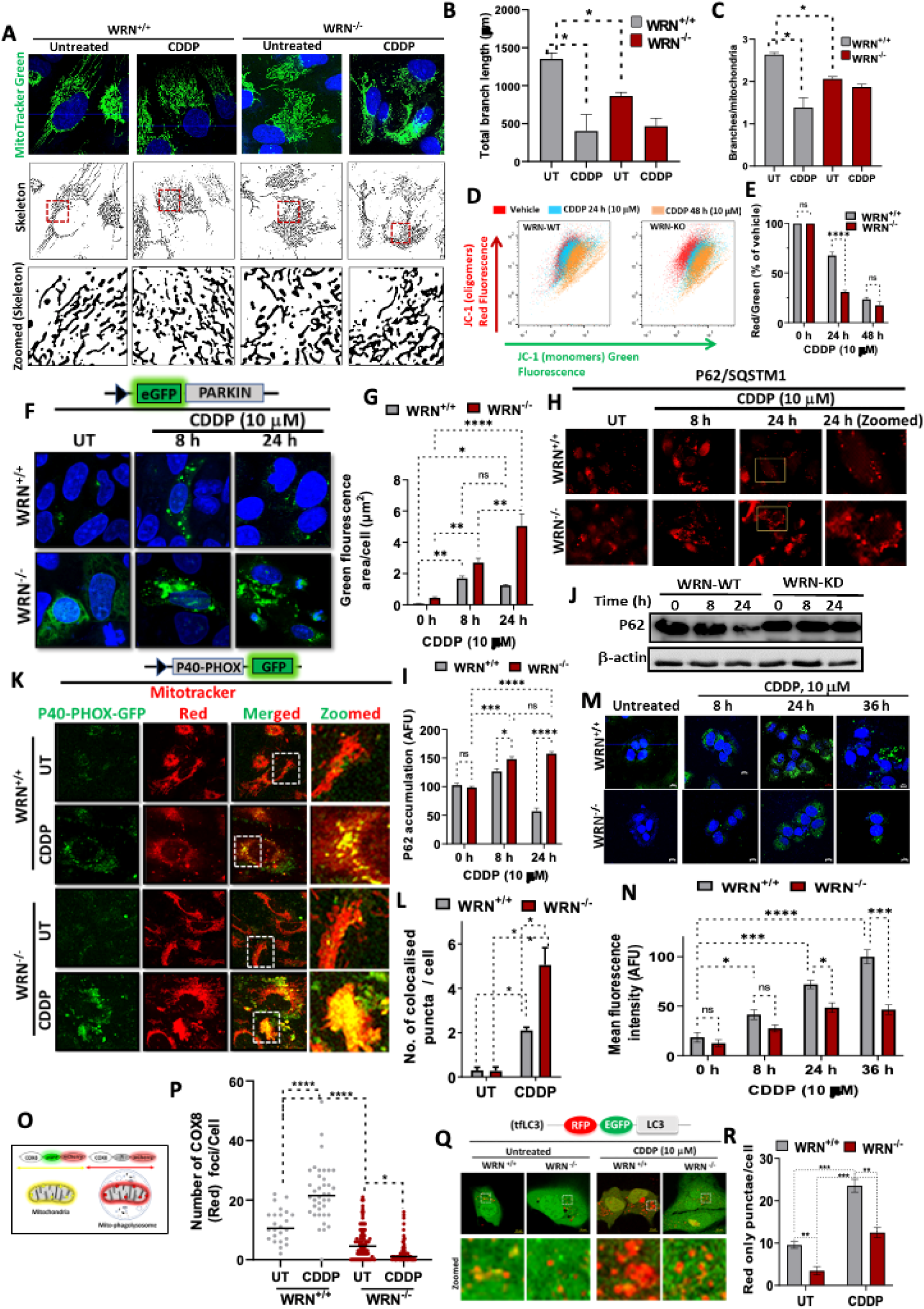
Mitochondrial dysfunction and defective mitophagy in WRN-deficient cells under cisplatin treatment. (A-C) Mitochondrial morphology in WRN^+/+^ and WRN^-/-^ U2-OS cells, treated with 10 µM cisplatin (CDDP) for 24 h, was visualized using Mito-Tracker Green (top) and skeletonized mitochondrial structures (middle and bottom). Zoomed-in images are shown on the bottom for detailed analysis. Quantifications were shown for total mitochondrial branch length (in µm) and branches per mitochondrion in untreated and cisplatin (CDDP)-treated WRN^+/+^ and WRN^-/-^ cells, indicating differences in mitochondrial network integrity. (D, E) Mitochondrial membrane potential was assessed and quantified by flow cytometry using JC-1 staining in WRN^+/+^ and WRN^-/-^cells treated with cisplatin (CDDP) for 24 or 48 h. The graph shows the JC-1 monomer fluorescence intensity, indicating changes in membrane potential. (F, G) Parkin translocation was observed by imaging eGFP-Parkin in WRN^+/+^ and WRN^-/-^ cells treated with cisplatin (CDDP) for 8 and 24 h. Nucleus were stained with DAPI (blue). Green fluorescence intensity of eGFP-Parkin was quantified and plotted. (H, I) P62/SQSTM1 accumulation was analyzed by immunofluorescence in WRN^+/+^ and WRN^-/-^ cells treated with cisplatin (CDDP, 10 µM) for 8 and 24 h. Zoomed-in images show the accumulation of p62 in the cytoplasm. P62 accumulation (in arbitrary fluorescence units, A.F.U.) in WRN^+/+^ and WRN^-/-^ cells, after cisplatin (CDDP) treatment, was quantified. (J) P62/SQSTM1 levels in WRN^+/+^ and WRN^-/-^ cells, after treatment with cisplatin (CDDP, 10 µM) for 0, 8, 24 h, was assessed. (K, L) Co-localization of P40-PHOX-GFP and MitoTracker Red was assessed by confocal imaging in WRN^+/+^ and WRN^-/-^ cells treated with cisplatin (CDDP, 10 µM) for 24 h. Co-localized puncta of P40-PHOX-GFP and mitochondria per cells, after treatment with cisplatin (CDDP, 10 µM) for 24 h in WRN^+/+^ and WRN^-/-^ cells was quantified. (M, N) WRN^+/+^ and WRN^-/-^ cells treated with cisplatin (CDDP, 10 µM) for 8, 24, and 36 h. Immunofluorescence of LC3 puncta was visualised by confocal microscopy and then quantified. (O, P) Schematic for detection of mitophagosomes and mitophagy flux using eGFP-mcherry-COX8 plasmid. Yellow fluorescence shows healthy mitochondria while red fluorescence shows mitophagosome fused with lysosome. The number of COX8 puncta (red only) per cell was quantified in untreated (UT) and cisplatin treated WRN^+/+^ and WRN^-/-^ cells. (Q, R) Representative images of WRN^+/+^ and WRN^-/-^ cells expressing tfLC3 (RFP-EGFP-LC3) under untreated and cisplatin (CDDP, 10 µM) conditions. Zoomed-in images highlight red-only puncta (autophagolysosomes) in WRN^-/-^ cells compared to WRN^+/+^ cells following cisplatin (CDDP) treatment. Red-only puncta per WRN^+/+^ and WRN^-/-^ cells, following cisplatin treatment, was quantified. All the values indicated are mean ± SEM (n = 3-4) \**p* <0.05, \*\**p* < 0.01, \*\*\**p* < 0.001 and \*\*\*\**p* < 0.0001.

Mechanistically, general mitophagy involves PINK1 stabilization on depolarized mitochondria, activation of Parkin, and subsequent ubiquitination of outer mitochondrial membrane proteins, which are recognized by p62/SQSTM1 and targeted to LC3-decorated autophagosomes for lysosomal degradation ^26^. To delineate which stage of this cascade is regulated by WRN, eGFP-Parkin expressing WRN^+/+^ and WRN^-/-^ cells were treated with cisplatin. WRN^+/+^ cells displayed a transient increase in Parkin puncta at 8 h that was resolved by 24 h, whereas WRN^-/-^ cells exhibited sustained Parkin accumulation at both time points (Fig. 2F, G). Similarly, WRN-deficient cells showed elevated levels of p62 puncta and protein content in response to cisplatin treatment as compared to control cells (Fig. 2H-J), suggesting that early recognition of damaged mitochondria is intact, but subsequent mitophagic processing is impaired. To evaluate mitophagophore formation, cells expressing the p40(phox)PX-eGFP probe were analyzed. The PX domain of p40(phox) is known to interact with phosphoinositide products of PI3Ks and thus used as a probe for measuring autophagosome assembly process around the depolarised mitochondria ^27^. Cisplatin induced greater p40(phox)PX puncta formation in WRN^-/-^ than WRN^+/+^ cells (Fig. 2K,L), indicating that initial mitophagophore assembly also proceeds normally in the absence of WRN. In contrast, LC3 puncta formation, representing mature autophagosomes, was robustly induced in WRN^+/+^ cells while both number and size of autophagosomes were reduced markedly in WRN^-/-^ cells (Fig. 2M,N), implicating a critical role of WRN in autophagosome maturation process. Further, a series of experiments were carried to validate the role of WRN in mitophagosome/autophagosome maturation. (1) Firstly, a specific traffic light mitophagy reporter construct was used, which consists of the COX8 mitochondrial targeting sequence placed N-terminal to an EGFP (pH-labile)-mCherry (pH-stable) fusion protein (Fig. 2O) ^28^. Cells expressing this construct show normal mitochondria in yellow (green + red fluorescence), whereas mitochondria within acidic compartments show red-only fluorescence, due to the selective quenching of EGFP fluorescence at low pH (Fig. 2O). At basal level, we detected significantly lower numbers of red-only-punctae in WRN^-/-^ cells *vs* WRN^+/+^ cells (Fig. 2O, P) while cisplatin treatment enhanced red-only-punctae in WRN^+/+^ cells while this was reduced in in WRN^-/-^ cells, respectively (Fig. 2O,P), confirming a role of WRN in mitophagosome maturation and their clearance. (2) Secondly, in order to rule out ectopic expression of COX8-EGFP-mCherry related anomaly in mitophagy, endogenous fusion of damaged mitochondria with lysosomes were assessed by mitochondria specific MitoTracker Green and lysosome specific LysoTracker Red dye. A similar pattern of lower level of basal and induced fusion associated mitophagosomes (yellow) were detected, validating a role of WRN in mitophagosome maturation and their clearance through mitophagy (Fig. S2G,H). (3) Thirdly, a different mitophagy inducing agent, CCCP (Carbonyl cyanide m-chlorophenyl hydrazone), was used. Further, a different tandem fluorescent-LC3 construct (tf-LC3: RFP (pH stable)-EGFP(pH labile)-LC3) was employed to monitor CCCP-induced formation of LC3-decorated-mitophagosomes (yellow punctae) and mitophagosome-lysosome fusion (mitophagolysosomes: Red-only punctae) (Fig. S2I). ^29^ Endogenous basal level of autophagolysosomes was found to be significantly lower in WRN^-/-^ cells *vs* WRN^+/+^ cells (Fig. S2I, J). CCCP treatment enhanced red-only-punctae in both WRN^+/+^ and WRN^-/-^ cells, but the induction was significantly compromised in WRN^-/-^ cells *vs* WRN^+/+^ cells (Fig. S2I, J). Moreover, the size of autophagolysosomes were distinctly smaller in WRN^-/-^ *vs* WRN^+/+^ cells. (4) Fourthly, a similar compromised autophagolysosomes, which also includes mitophagolysosomes, were detected in response to cisplatin treatment also (Fig. 2Q, R). (5) Fifthly, we sought to know whether the defective autophagosome maturation in WRN^-/-^ cells is associated with cisplatin/CCCP specific treatment or a general phenomenon. To this end, we assessed autophagolysosomes (red only tf-LC3 punctae) formation in response to amino acid starvation, which is known to rapidly induce autophagy process ^30^. We detected severely compromised autophagolysosomes formation, in term of their both number and size, in WRN^-/-^ cells than WRN^+/+^ cells (Fig. S2K, L). Collectively, these comprehensive analyses demonstrate that WRN plays a pivotal role in maturation of autophagosomes and mitophagosomes, ensuring efficient clearance of damaged mitochondria and the maintenance of mitochondrial quality control under both basal and stress conditions.

### WRN helicase regulates ATG7 expression and autophagosome maturation *via* G4-DNA resolution and U2AF35-dependent mRNA processing

Our findings raised an intriguing mechanistic question regarding how WRN, a predominantly nuclear RecQ helicase, influences the cytosolic process of autophagosome maturation. Among the core autophagy-related proteins (Beclin-1, ATG3, ATG5, ATG7, ATG12, etc.; Fig. 3A), both basal and cisplatin (CDDP)-induced expression of ATG7 were markedly reduced in WRN^-/-^ cells compared to WRN^+/+^ counterparts (Fig. 3B; Fig. S3A-F). ATG7, an E1-like activating enzyme, is indispensable for autophagosome biogenesis, as it mediates the conjugation of LC3-I with phosphatidylethanolamine (PE) to generate lipidated LC3-II ^31^. Consistent with reduced ATG7 expression, both basal and cisplatin-induced levels of LC3-II were significantly low in WRN-deficient cells relative to controls (Fig. 3B, Fig. S3F). To exclude the possibility that lower LC3-II levels resulted from enhanced autophagic flux, LC3-II accumulation was analyzed following bafilomycin A1 (BafA1) treatment. In agreement with our earlier observations, LC3-II level remained markedly reduced, accompanied by higher p62/SQSTM1 accumulation in WRN^-/-^ cells relative to WRN^+/+^ controls (Fig. S3G), confirming a defect in LC3-II–mediated autophagosome assembly. Furthermore, siRNA-mediated depletion of ATG7 phenocopied WRN loss, leading to increased mitochondrial mass, shortened branch length, and diminished network connectivity (Fig. S3H-J), recapitulating the defective mitophagy phenotype of WRN-deficient cells. Collectively, these findings establish WRN as a critical regulator of ATG7 expression and autophagosome maturation.

**Figure 3:**
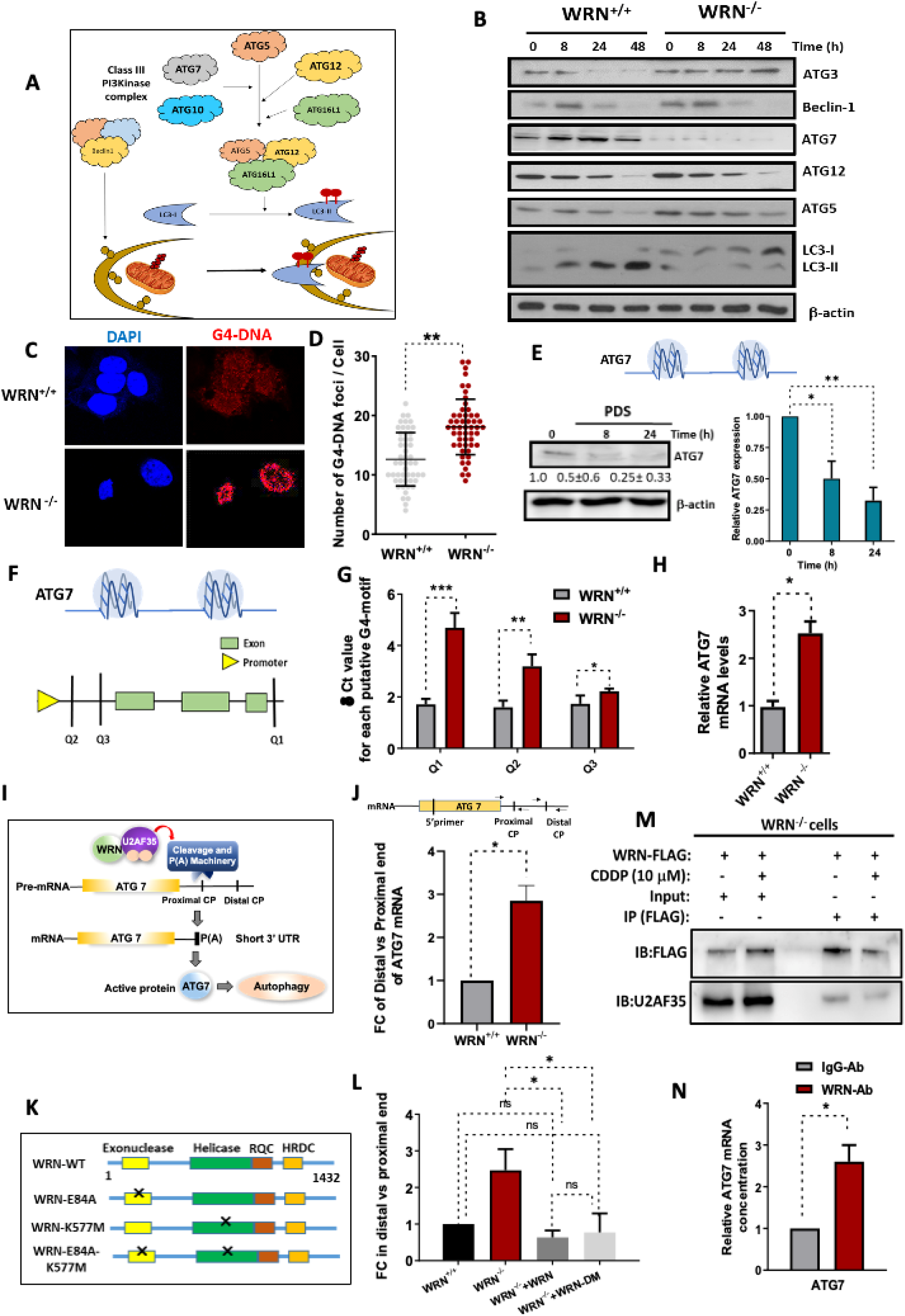
Mitophagy, G4 quadruplex and pre-mRNA maturation in WRN-deficient cells. (A) Schematic representation of the autophagy pathway, highlighting key proteins such as ATG7, ATG5, Beclin-1 and Class III PI3K complex involved in the lipidation of LC3 and formation of autophagosome. (B) Western blot analysis of autophagy-related proteins in WRN^+/+^ and WRN^-/-^ cells treated with cisplatin (CDDP, 10 µM) for indicated time. Proteins probed are ATG3, Beclin-1, ATG7, ATG12, ATG5, and LC3-I/II. β-actin was used as a loading control. (C, D) Immunofluorescence of G4-DNA (red) in WRN^+/+^ and WRN^-/-^ cells, co-stained with DAPI (blue). Quantification of G4-DNA foci per cell in WRN^+/+^ and WRN-/- cells were plotted (E) ATG7 protein expression in WRN^+/+^ and WRN^-/-^ cells treated with Pyridostatin (PDS: a G4 stabilizer) for indicated time interval. Protein levels were measured by Western blotting. (F) Graphical schematic of ATG7 gene showing potential G-quadruplex-forming regions (Q1, Q2, Q3) with respect to promoter and intron/exon. (G) Chromatin immunoprecipitation with G4 antibody was performed in WRN^+/+^ and WRN^-/-^ cells. G4-DNA was detected and quantified with the RT-PCR primer designed at Q1, Q2 and Q3 putative site in ATG7 gene. (H) Relative total cellular ATG7 mRNA levels in WRN^+/+^ and WRN^-/-^ cells was determined by qRT-PCR. (I) Schematic model illustrating the role of U2AF35 and probable involvement of WRN in the regulation of nascent ATG7 mRNA processing. The process involves cleavage at the proximal and distal cleavage/polyadenylation (CP) sites, leading to the formation of a shorter and mature 3’-UTR for the expression of active ATG7 protein, which subsequently activates autophagy. (J) Quantification of ATG7 mRNA processing in WRN^+/+^ and WRN^-/-^ cells. The fold change (FC) of distal *versus* proximal end ATG7 mRNA between the two cell types was analyzed and quantified. (K) WRN mutant constructs used in the study. Schematic representation of WRN^+/+^ and WRN mutants, highlighting the functional domains (Exonuclease, Helicase, RQC, HRDC) and the sites of mutations for generation of helicase-dead (K577M) and/or exonuclease-dead (E84A) WRN proteins. (L) ATG7 mRNA (distal and proximal) in WRN^-/-^ cells and WRN^-/-^ cells complemented with WRN-WT and mutant, was analyzed and quantified for the fold change of distal *versus* proximal end ATG7 mRNA. (M, N) Co-immunoprecipitation of FLAG-tagged-WRN with U2AF35 in WRN^-/-^ cells. Cells were treated with cisplatin (CDDP, 10 µM) for 24 h, and lysates were immunoprecipitated using an IgG control or anti-FLAG antibody. The FLAG and U2AF35 proteins were detected by immunoblotting (IB). (N) RNA immunoprecipitation (RIP) was performed with anti-WRN antibody (WRN-Ab) or IgG antibody (IgG-Ab), and the qRT-PCR was done with ATG7 primer. Relative ATG7 mRNA levels in RNA pull down with anti-WRN antibody (WRN-Ab) or IgG antibody (IgG-Ab) treatment was plotted. All the values indicated are mean ± SEM (n = 3-4), ns denotes nonsignificant, \**p*<0.05, \*\**p* < 0.01 and \*\*\**p* < 0.001.

Transcriptional regulation of numerous genes is influenced by the presence of G-quadruplex (G4) DNA structures within their regulatory or coding regions ^32,33^ WRN helicase has been shown to unwind and resolve G4 structures, thereby facilitating gene expression and genome stability ^34,35^. Based on our results on ATG7 protein expression (Fig. S3B), we hypothesized that WRN may regulate ATG7 transcription by resolving G4-DNA structures within the ATG7 locus. We observed a significantly higher level of cellular G4-DNA, as detected by a G4-specific antibody, in WRN^-/-^ cells relative to WRN^+/+^ cells (Fig. 3C, D). To examine whether stabilization of G4-structures impacts ATG7 expression, we assessed ATG7 protein levels following Pyridostatin (PDS), a selective G4-DNA stabilizing ligand ^36^, treatment. Notably, PDS exposure led to a pronounced reduction in ATG7 protein levels in U2-OS cells (Fig. 3E), indicating that G4-DNA stabilization negatively regulates ATG7 expression. To identify potential G4-forming motifs within the human ATG7 gene, we performed *in-silico* G4 prediction using the QGRS Mapper tool (http://bioinformatics.ramapo.edu/QGRS/index.php) ^37^. Three high-confidence G4 motifs (designated Q1, Q2, and Q3) were identified, with Q2 and Q3 located within 5’UTR (untranslated region) and Q1 within 3’UTR region of ATG7 gene. Chromatin immunoprecipitation (ChIP) using a G4-specific antibody, followed by qPCR analysis, revealed significantly enriched occupancy of all three G4 motifs in ATG7 gene in WRN^-/-^ cells compared to WRN^+/+^ cells (Fig. 3G, H), with strong signal at Q1 site suggesting that WRN facilitates ATG7 expression by resolving G4-DNA structures within its gene locus. Unexpectedly, despite reduced ATG7 protein levels, ATG7 mRNA expression was strikingly higher in WRN^-/-^ cells relative to wild-type cells (Fig. 3H), implying a potential post-transcriptional or mRNA maturation defect associated with WRN loss. Notably, Michael Green group previously reported an elegant finding that U2AF35, a core subunit of the U2 auxiliary factor (U2AF) complex, is essential for processing ATG7 pre-mRNA into its mature transcript ^38^. U2AF35 recruits the cleavage and polyadenylation (CPA) machinery to catalyze endonucleolytic cleavage and poly(A)-tail addition at the proximal 3′-UTR cleavage/polyadenylation (CP) site (Fig. 3I) ^38,39^. A U2AF35(S34F) mutant variant promotes aberrant processing at a distal CP site, generating a longer, inefficiently translated ATG7 transcript (Fig. 3I) ^38^. Consequently, reduced ATG7 expression impairs autophagy and promotes genomic instability and malignant transformation ^38,40^. Given these insights, we investigated whether WRN modulates U2AF35-mediated ATG7 mRNA processing. Since our previous qPCR assay for ATG7 mRNA expression (Fig. 3H) did not discriminate between nascent pre-mRNA and mature mRNA, we further used primers targeting the distal CP site (long 3′-UTR) and proximal CP site (short 3′-UTR) of ATG7 (Fig. 3J; Upper panel). Strikingly, the ratio of distal-to-proximal ATG7 transcripts was approximately threefold higher in WRN^-/-^ cells relative to WRN^+/+^ cells (Fig. 3J), consistent with defective post-transcriptional processing in WRN-deficient cells. Considering that WRN can also resolve RNA secondary structures, it is plausible that its helicase and/or exonuclease activities may cooperate with U2AF35 to facilitate accurate ATG7 mRNA maturation. Complementation of WRN^-/-^ cells with wild-type WRN completely restored ATG7 mRNA processing, as evidenced by the reduction of distal-end transcripts (Fig. 3K, L), confirming a role for WRN in generating mature ATG7 mRNA with a short 3′-UTR. Interestingly, complementation of WRN^-/-^ cells with a helicase/exonuclease double mutant (WRN-DM; K577M/E84A) similarly rescued the phenotype (Fig. 3L,M), suggesting that WRN promotes ATG7 pre-mRNA maturation *via* its non-enzymatic mechanism. To test whether WRN directly associates with U2AF35 during ATG7 mRNA processing, two complementary assays were performed. (1) Immunoprecipitation (IP) of WRN-FLAG from complemented WRN^-/-^ cells demonstrated a robust interaction with U2AF35 under both basal and cisplatin-treated conditions (Fig. 3M). (2) RNA immunoprecipitation (RIP) using anti-WRN antibody followed by qPCR revealed significant enrichment of ATG7 mRNA, compared with IgG controls (Fig. 3N). Together, these findings establish that WRN directly interacts with U2AF35 to promote post-transcriptional processing of ATG7 pre-mRNA, thereby coupling nuclear RNA maturation with cytosolic autophagy regulation. Collectively, our findings establish that WRN facilitates cellular autophagy by resolving G4-quadruplex structures within the ATG7 gene and promoting U2AF35-dependent post-transcriptional processing of nascent ATG7 mRNA.

### WRN suppresses R-loop generation at ATG7 gene by resolving G4-quadruplex motifs in cancers

Monnat *et al* previously demonstrated that the WRN helicase functions as a G-quadruplex (G4)–resolving enzyme at numerous chromosomal loci, thereby influencing transcriptional regulation and gene expression ^41^. More recently, Franchitto and Pichierri *et al* provided compelling evidence that WRN mitigates transcription–replication conflicts and suppresses the accumulation of R-loops (DNA-RNA hybrids) in human cells ^42^. Building upon these findings, our results suggest an additional mechanistic role for WRN may coordinate G4-DNA unwinding associated prevention of R-loop formation, thereby safeguarding transcriptional fidelity. To elucidate this mechanism, we focused on the ATG7 gene as a representative candidate, given its pronounced WRN-dependent regulation observed in our study. Immunofluorescence analysis using the R-loop–specific antibody S9.6 revealed significantly elevated basal R-loop levels in WRN^-/-^cells compared with WRN^+/+^ counterparts (Fig. 4A). Upon cisplatin treatment, R-loops were markedly increased in WRN^+/+^ cells, whereas this induction was largely attenuated in WRN ^/^ cells (Fig. 4A,B). Consistent with these findings, dot-blot analysis of chromatin-associated R-loops confirmed robust basal accumulation of R-loops in WRN-deficient cells, Moreover, cisplatin treatment enhanced R-loops formation several folds in WRN^+/+^ cells while only a minor enhancement was observed in WRN^-/-^ cells (Fig. 4C,D). Signals in dot blot were abolished by RNase H treatment, confirming antibody specificity for R-loop. Given that our G4-DNA ChIP analysis revealed higher enrichment of three distinct G4 motifs (Q1, Q2 & Q3) within the ATG7 locus in WRN ^/^ cells relative to wild-type (Fig. 3H), it is plausible that unresolved G4 structures may hinder transcriptional elongation, leading to the accumulation of ATG7 pre-mRNA as R-loop intermediates. In this regard, chromatin-bound RNA analysis demonstrated more than a two-fold increase in ATG7 pre-mRNA containing the distal 3′-UTR as well as ORF region in the form of R-loops in WRN^-/-^ cells compared to WRN^+/+^ cells (Fig. 4E, F), substantiating a direct role for WRN in resolving G4-DNA–associated R-loop structures within the ATG7 gene. These findings suggest that impaired resolution of G4-DNA and R-loops and defective nascent ATG7 mRNA maturation process in WRN-deficient cells compromises ATG7 expression, leading to defective autophagy and mitophagy in cancer cells.

**Figure 4:**
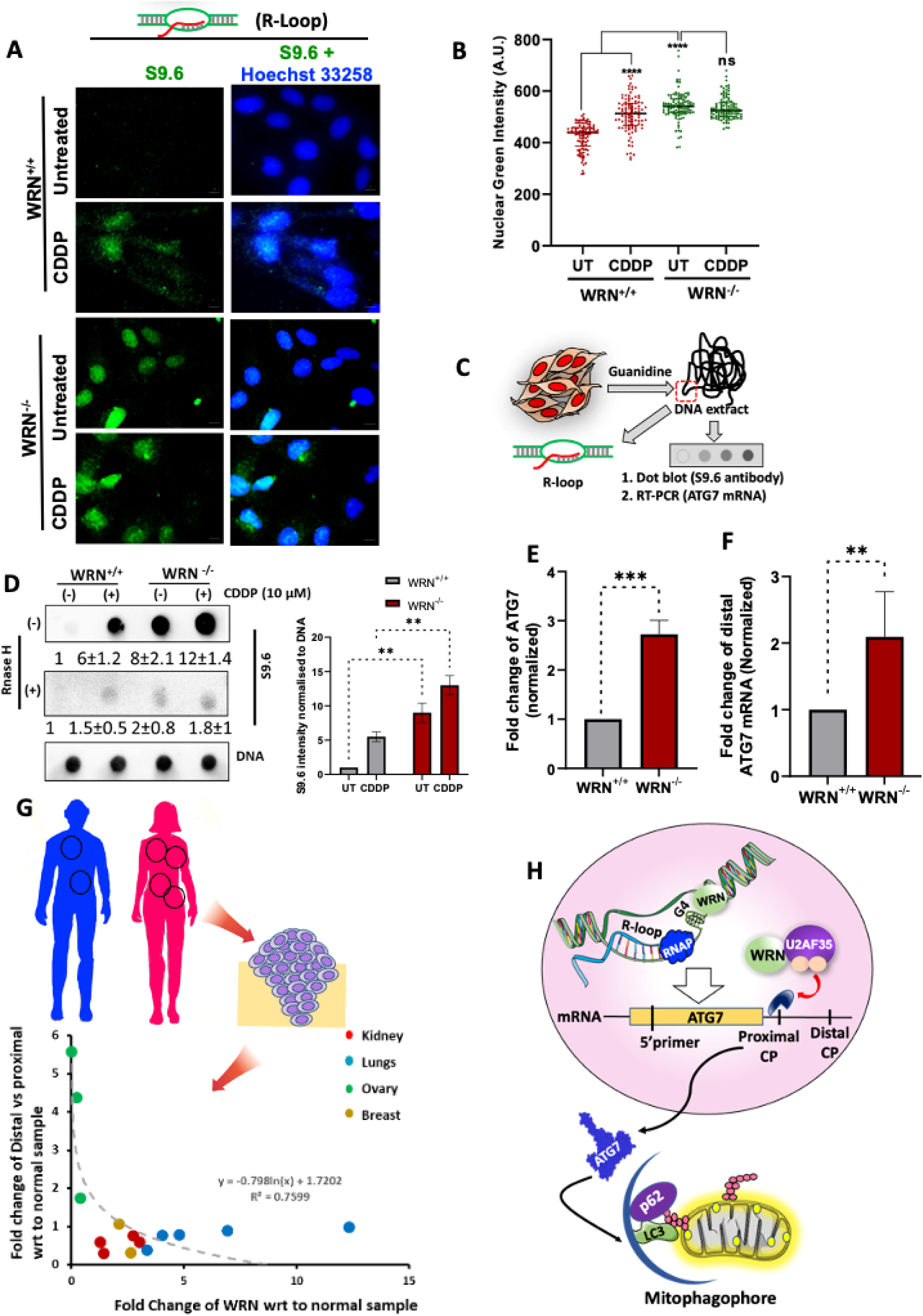
Role of WRN in R-loop formation during ATG7 mRNA maturation and processing. (A, B) R-loops in WRN^+/+^ and WRN^-/-^ cells was detected, in untreated and CDDP-treatment (10 µM, 24 h), by immunofluorescence microscopy using the S9.6 antibody. Nuclear R-loops was quantified in WRN^+/+^ and WRN^-/-^ cells. (C, D) Graphical representation for preparation cellular R-loop and detection by slot blot. R-loops samples were prepared from untreated and cisplatin (CDDP, 10 µM) treated-cells and R-loops was analyzed by dot blots. In control, R-loops lysates were treated with RNase H (+) to confirm the R-loop specificity. (E, F). Chromatin bound RNA was isolated from WRN^+/+^ and WRN^-/-^ cells. ATG7 mRNA (proximal and distal) was quantified by RT-PCR. (G) Analysis of mRNA status for ATG7 mRNA (proximal vs distal) and WRN across different cancer tissues (kidney, lungs, ovary and breast), derived from cancer patients. The graph plots the fold change of distal *versus* proximal ATG7 mRNA and WRN. A regression analysis is shown to highlight the relationship between WRN expression and ATG7 processing in different tissues. (H) Proposed mechanism for the regulation of WRN-mediated autophagy/mitophagy. WRN resolves G4-DNA in ATG7 gene. Moreover, WRN in cooperation with U2AF35, regulates ATG7 mRNA processing, affecting R-loop formation and ultimately promoting mitophagy through autophagic machinery, involving p62, LC3, and the formation of mitophagosomes. All the values indicated are mean ± SEM (n = 3-4), ns denotes non-significant, \*\**p* < 0.001, \*\*\**p* < 0.001 and \*\*\*\**p* < 0.0001.

Finally, to evaluate the clinical relevance of WRN deficiency and its association with ATG7 dysregulation, we examined mRNA derived from tumor tissue samples from patients with lung, breast, ovarian, and kidney cancers. Quantitative PCR analysis revealed a strong inverse correlation between WRN mRNA levels and ATG7 transcripts (fold change of distal *vs* proximal mRNA) (Fig. 4G), supporting the notion that WRN loss is associated with defective ATG7 mRNA maturation and impaired autophagic competence in tumors. Given the emerging clinical significance of WRN as a synthetic-lethal target and therapeutic vulnerability in multiple cancers ^43^, our study provides mechanistic insight into how WRN deficiency or pharmacological targeting of WRN may influence therapeutic outcomes through dysregulated autophagy/mitophagy pathways in cancers.

## Discussion

Multiple independent large-scale studies employing RNA interference and CRISPR-Cas9 mediated gene editing have demonstrated that loss of WRN function results in selective or synthetic lethality in cancer cells exhibiting microsatellite instability (MSI), a consequence of defective DNA mismatch repair (MMR) pathways ^14,15,44,45^. Two comprehensive efforts-Project DRIVE (Novartis) ^46^ and Project Achilles (Broad Institute)^47^-systematically interrogated genome-wide dependencies across extensive panels of human cancer cell lines representing diverse tissue origins. These large-scale functional genomic screens identified multiple synthetic lethal interactions among curated gene sets and converged on PRMT5 and WRN as key, high-confidence druggable vulnerabilities. Building upon these findings, pharmacological inhibitors of WRN are currently under clinical evaluation as potential therapeutic agents for MSI-high malignancies^17,48^. Despite this progress, the functional consequences of WRN inhibition or deficiency beyond its canonical role in the DNA damage response (DDR) remain incompletely understood. In particular, the contribution of WRN to mitophagy and autophagy regulation in cancer is poorly characterized. In *C. elegans* invertebrate model, Bohr and his coworkers has showed that WRN regulates mitophagy through ULK1-BNIP3L axis ^18^. In cancer, however, association of WRN deficiency with mitophagy has not been explored yet. Elucidating the molecular mechanisms linking WRN to mitophagy may provide critical insights into the cellular stress responses underlying cancer cell survival and inform therapeutic strategies targeting WRN-deficient or WRN-inhibited tumors.

G-quadruplexes are enriched in regulatory locations of several key genes such as promoters, untranslated regions and gene bodies, and hence influences their expressions^49^. G4-DNAs on the non template or template DNA strand may modulate polymerase II progression, leading to transcriptional blockage and R-loop accumulation ^50^. Cells deploy G4-resolving helicases, e.g., WRN, BLM, FANCJ, DHX36, to prevent/resolve G4s to restore transcriptional continuity ^51^. In WS patient fibroblasts, significant enrichment of G4 motifs are found, especially in downregulated genes ^41^. However, precise role of WRN on resolving G4s in autophagy genes and its impact on functional autophagy process in cancers are not known. In the current investigation, heavy downregulation of ATG7 protein, among a battery of autophagy regulating proteins, was found at endogenous level and in response to cisplatin in WRN-deficient cancer. In association with ATG7 downregulation, mitophagosomes and autophagosomes formation are also severely down regulated in WRN-deficient cancer. Considering the profound role of ATG7 in autophagy, we focused on unravelling the hitherto unknown role of WRN in resolving G4-motifs, if any, in ATG7 gene expression. To this end, we identified three high-confidence G4 motifs in 5′-UTR and 3′-UTR of ATG7 gene, using G4 prediction tool (the QGRS Mapper). Further, ChIP assay using a G4-DNA specific antibody, followed by qPCR analysis, revealed significantly enriched occupancy of all three G4 motifs in ATG7 gene in WRN^-/-^ cells compared to WRN^+/+^ cells (Fig. 3G, H), suggesting for the first time that WRN may facilitates ATG7 expression by resolving G4-DNA structures within its gene locus (Fig. 4H). However, when we assessed ATG7 mRNA level, it was found to be unexpectedly higher in WRN^-/-^ cells than WRN^+/+^ cells. In both pyridostatin (G4 stabilizing agent) treated cells and WRN^-/-^ cells, ATG7 expression was significantly lower, which cannot be explained with higher mRNA level in WRN^-/-^ cells but indicated a compromised post-transcriptional processing of ATG7 mRNA in WRN-deficient cells. A recent elegant report has shown that U2AF35 in association with cleavage and polyadenylation (CPA) machinery cleaves nascent ATG7 mRNA to generate the mature ATG7 for its efficient translation. However, whether this process is modulated by WRN is not yet known. We demonstrate for the first time that WRN physically associates with the splicing factor U2AF35 to facilitate the post-transcriptional processing of ATG7 pre-mRNA, in a non-enzymatic manner, thereby establishing a mechanistic link between nuclear RNA maturation and cytoplasmic autophagy regulation (Fig. 4H). Several reports show U2AF35 mutations change splicing/3′ end maturation of mRNA and the downstream oncogenic consequences and defective autophagy, including ATG7 loss ^38,52^, leading to initiation of malignant transformation. In this connection, our findings show a novel context of WRN loss in defective autophagy/mitophagy, in addition to its known association with genome instability, which may further contribute to cancer disposition characteristic in WS patients and initiation of malignant transformation in various tissue types with WRN-deficiency. In addition to differential mRNA splicing regulation, high levels of G4-DNA in WRN^-/-^ cells may influence R-loop accumulation and gene expression in WRN-deficient cells. Previously, association of R-loops in ATG7 gene was reported in UV-irradiated skin cells ^53^. In the current investigation, we show that WRN regulates R-loops abundance (ATG7 mRNA) in ATG7 gene, influencing the U2AF35 mediated processing of ATG7 distal mRNA (Fig. 4H). It has been validated in clinical setting also, as an inverse relation of WRN and ATG7 (distal vs proximal) mRNA expression were observed.

In summary, impairment of autophagy and mitophagy in WRN-deficient cancer cells profoundly disrupts mitochondrial homeostasis, leading to reduced oxidative phosphorylation, elevated ROS accumulation, and broad metabolic dysregulation that can influence therapeutic responsiveness. Such defects in mitochondrial quality-control pathways are likely to exacerbate malignant progression in the WRN-deficient context. Collectively, our findings identify WRN as a critical upstream regulator of autophagy and mitophagy, essential for sustaining mitochondrial dynamics and metabolic integrity in tumor cells (Fig. 4H). Given that ATG7 has been implicated as a suppressor of tumor progression and metastasis ^54^, WRN-deficiency-driven attenuation of ATG7 expression may contribute mechanistically to the observed oncopathogenic phenotype. Comprehensive, genome-wide multi-omics profiling of R-loops and G-quadruplex DNA structures in WRN-deficient cancers will further clarify the mechanistic crosstalk between defective autophagy pathways and compromised genome maintenance. Such studies may ultimately refine the therapeutic rationale for targeting WRN-associated vulnerabilities in cancer.

## Materials and Methods

### Materials

Antibodies for BECN1 (#3495P), LC3A/B I/II (#12741), LC3B (#3868P), ATG5 (#12994P), ATG7 (#8558P), ATG3(#3145P), ATG12 (#4180) were procured from Cell Signaling Technology (Danvers, MA, USA). Antibodies against p62, WRN (#SC5629) and CRISPR-Cas9 double nickase plasmids (control and WRN) were from Santa Cruz biotechnology (Santa Cruz, CA). Rabbit Anti-DNA G4 Quadruplex antibody (MABE1126), mouse monoclonal anti-DNA-RNA hybrid antibody (#MABE1095), mouse monoclonal anti-DNA antibody (#MAB030), rabbit monoclonal β-actin (A5316), mouse monoclonal anti-Flag (F1804) and mouse monoclonal anti poly-ADP-ribose antibody (#MABC547) were purchased from Sigma Aldrich (St Louis, MO, USA). Rabbit polyclonal anti-U2AF35 from Bethyl Laboratories. MitoTracker Green, LysoTracker Red, Lipofectamine 2000, Prolong anti-fade gold, Alexa Flour-488/555/595 (#A21123/#A31570), JC-1, NBD-glucose and DCF-DA fluorescent dye from Invitrogen (Life Technologies, Carlsbad, CA). All other reagents like Bafilomycin, Pyridostatin and Cisplatin were obtained from Sigma Chemicals (St. Louis, MO), unless mentioned in the respective places.

### Cell Culture

U2-OS (human osteosarcoma) cells was procured from ECACC and cultured in Dulbecco’s Modified Eagle Medium supplemented with 10% fetal bovine serum and Penicillin-Streptomycin-Amphotericin B cocktail. Cells were maintained at 37 under 95 % relative humidity and 5% CO_2_.

### Generation of WRN knockout cells

U2-OS cells were transfected with WRN CRISPR-Cas9 and control CRISPR-Cas9 double nickase plasmid system (SCBT) and followed the manufacturer’s protocol. Cells were selected with puromycin for stable knockout (WRN^-/-^) and then confirmed by immunoblotting.

### Measurement of mitochondrial respiration rate

For determining oxygen consumption rate (OCR), 1 × 10^5^ cells/well were seeded in 24 well respirometric analysis plates. After 24 h of serum starvation, the OCR measurement was done using an XF–24 extracellular flux analyser (Seahorse Bioscience, MA, USA) using Seahorse XF Mito Stress Test Kit and analysed using WAVE software.

### NBD glucose uptake assay

WRN^+/+^ and WRN^-/-^ cells (0.5 x 10^5^ cells/well), seeded in 6-well plate for 24 h, washed with PBS, and 100 µM of 2-N-7-(nitrobenz-2-oxa-1,3-diazol-4-yl)amino-2-deoxy-d-glucose (2-NBDG) was added in starved condition at 37 °C for 1h. The cells were trypsinised, washed with PBS and equal number of cells were analysed for the glucose uptake by measuring the fluorescence of NBDG with a Partec CyFlow® Space flow cytometer by using the FlowJo software.

### Immunoblotting

Western blotting was performed as previously reported ^55^. Protein samples were prepared by adding TNN buffer (20 mM Tris (pH 7.4), 250 mM NaCl, 0.05% NP40) that contained protease and phosphatase inhibitor cocktail. After protein quantification by Bradford, equal amount of protein was loaded on SDS-PAGE gel and then transferred the proteins to nitrocellulose membrane. The nitrocellulose membrane was probed with appropriate (primary and secondary) antibody after blocking with skimmed milk. The membrane was developed by adding substrate for HRP and detected chemi-luminescence. Protein amounts (arbitrary unit, mean ± S.D) were quantified by density scanning and normalized by considering that of untreated/vehicle treated sample as 1 and normalizing with loading control.

### ATP assay

The cells (1 x 10^5^ /well) were seeded in 6-well plates and treated with vehicle (0.1% DMSO) or cisplatin (10 μM) at 37 °C for 24 h, cells were washed with PBS, lysed in TNN buffer. ATP level was quantified as per the manufacture protocol of ATP Determination Kit (A22066 from Invitrogen) and measured the luminescence (RFU) per 10 μg protein by microplate reader from BMG Labtech.

### JC1 staining

WRN^+/+^ and WRN^-/-^ cells (1×10^5^ cells/well), seeded in 6-well plate were treated with cisplatin for 24 h, washed with PBS, and incubated with JC-1 (20 μM) for 1 h. The cells were trypsinised, washed with PBS and equal number of cells were analysed for the mitochondrial potential loss from the shift of emission from green (∼525 nm) to red (∼590 nm) with a Partec CyFlow Space flow cytometer. Data analysis was carried out using the FlowJo Software (FlowJo V10 Version v10.7.1. Ashland).

### MitoTracker and LysoTracker staining

WRN^+/+^, WRN^-/-^, ATG7-WT and ATG7-KD (1×10^5^ cells/well) seeded in 6-well plate, were treated with cisplatin for 24 h, washed with PBS, and incubated with MitoTracker Green (200 nM) and LysoTracker Red (200 nM) for 30 min. The cells were trypsinized, washed with PBS and equal number of cells was analyzed with a Partec CyFlow Space flow cytometer by sing the FlowJo software. For microscopic staining, cells were seeded on coverslip and stained with 200 nM of MitoTracker Green and LysoTracker Red for 30 min and visualised under fluorescent microscope (LSM-780) and then analyses of mitochondrial network were performed by Fiji software.

### DCF-DA and mitoROS assay

WRN^+/+^ and WRN^-/-^ cells (1 x 10^5^ cells/well), seeded in 6-well plate for overnight and treated with cisplatin (10 µM) for 24 h, washed with PBS, and incubated with 2’,7’-dichlorofluorescin diacetate (DCF-DA) for 40 min. The cells were trypsinised, washed with PBS and equal number of cells were analysed for the DCF fluorescence with a Partec CyFlow Space flow cytometer by using the FlowJo software. In order to measure intra-mitochondrial superoxide/reactive oxygen species, 3 x 10^5^ cells/well were seeded in a 6-well plate. Cells were treated with MitoSOX agent (5 μM), in 37 °C incubator, protected from light, for 30 min followed by CDDP (10 µM) treatment for 0, 24 and 48 h. Fluorescence was measured using flow cytometry in the red channel and fold change is plotted as in bar graph.

### Plasmids and transfection

Cells were transfected with plasmids of pEGFP-Parkin (Addgene #45875), pEGFP-LC3 (Addgene #24920) and mRFP-EGFP-LC3 (tf-LC3; Addgene #21074), P(40)PX-EGFP (Addgene #19010), COX8-EGFP-mCherry (Addgene #78520), Flag-WRN ^13^, respectively. These plasmids were transfected transiently by using Lipofectamine 2000 reagent in WRN^+/+^ and WRN^-/-^ cells (1 x 10^5^ cells/well) and corresponding empty backbone vector was used for control experiment. After 24 h, the expression was confirmed by fluorescence microscope.

### Fluorescence microscopy

Cells expressing EGFP-Parkin, P(40)PX-EGFP, EGFP-LC3, mRFP-EGFP-LC3 (tf-LC3) and tf-COX8 were treated with cisplatin (10 μM) for indicated time periods. The images for the cells with requisite fluorescence were captured by Zeiss LSM780. Specific fluorescence intensity or fluorescent puncta per cell was assessed manually from at least 3 different experiments and at least 50-100 cells per group were scored in each experiment.

### Site directed mutagenesis

WRN double mutants (K577M and E84A) were generated as previously described ^13^, using “QuikChange-II” site-directed mutagenesis kit (Stratagene, CA, USA) as per manufacturer’s protocol and the mutations were confirmed by sequencing.

### Immunofluorescence assay

The cells (1 x 10^5^ /well) were seeded on coverslip in 6-well plates and treated with vehicle (0.1% DMSO) or cisplatin (10 μM) at 37 °C for indicated time points, cells were washed with PBS, fixed with 1% PFA (3 min), permeabilized in Triton X-100 (0.5% in PBS) on ice for 10 min, washed with PBS, and subjected to immunofluorescence using antibody against p62, G4-DNAs and LC3and appropriate secondary antibody with indicated Alexa probe. Slides were mounted in Prolong Gold anti-fade reagent with DAPI. For G4 quadruplex immunofluorescence, Cells were fixed in 4% paraformaldehyde (PFA, Sigma) in PBS and then permeabilized with 0.5% Tween 20 in PBS. Cells were treated with 20 mg/500 ml RNase A(Invitrogen) and subsequently blocked in 5% normal bovine serum (Calbiochem) for 12 h at 4 °C. Cells were then incubated with purified G4-DNA antibody for overnight at 4 °C. Cells were washed with PBST and then incubated with Alexa Fluor 596 goat anti-mouse antibody (Invitrogen) followed by staining for DNA and mounted coverslips with Prolong gold antifade (Invitrogen). Images were captured using an LSM 780 Meta laser scanning confocal microscope.

### G4-quadruplex mapping

The QGRS mapper (http://bioinformatics.ramapo.edu/QGRS/index.php) was used to identify the putative G4-DNA quadruplex structures in ATG7 gene. Four putative G4 quadruplex sites were taken further for DNA ChIP analyses ^37^.

### G4-DNA ChIP

WRN^+/+^ and WRN^-/-^ cells (1 x 10^7^ cells per flask) were seeded in two tissue culture flasks for 24 h and fixed with 1% paraformaldehyde for 10 min at RT. Cross-linking reaction was stopped by adding glycine to a final concentration of ∼0.12 M, followed by incubation for 10 min at RT. Cells were washed twice with 10 ml of ice-cold PBS. Cells were scraped and the pellet was lysed with hypotonic buffer. The pellet was used as nuclear suspension followed by sonication. Sonicated mixture was treated with RNase A (10 µg/ml) at 37 °C for 20 min. Further, the mixture was treated with proteinase K (40 µg/ml) for 2 h at 65 °C. Chromatin fragment size distribution was analysed by using 2% (wt/vol) agarose gel electrophoresis. With fragment distribution of 100–500 bp, 2000 ng/sample were subjected to ChIP by following a protocol as reported previously ^56^.

### qRT-PCR assay

Isolation of total RNA was performed using TRIzol reagent (Ambion, Life technologies). Two microgram of total RNA from each sample was used to generate cDNA by using TOPscript™ cDNA Synthesis Kit (Cat. No. EZ005M, Life technology). The specific primers (mentioned in Table1) were used to amplify in real-time reaction and monitored using SYBR green (from Biorad) in CFX96 Touch Real-Time PCR System (Biorad, US). The Ct values obtained for the desired amplicon were normalized to control gene GAPDH expression. But for calculation of fold change of distal vs proximal mRNA of ATG7, the distal amplicon was normalized with respective proximal.

### R-loop (RNA/DNA hybrid) immunofluorescence

RNA-DNA hybrids was detected microscopically by following the protocol described previously with minor modifications. Cells were seeded on coverslips overnight, then treated with 10 µM cisplatin for 24 h. Following treatment, cells were fixed with 4% paraformaldehyde in PBS for 10 min, then permeabilized with 0.5% Triton X-100 (ice-cold). RNase A digestion was performed using a cocktail containing 6 µg/mL RNase A, 10 mM Tris-HCl (pH 7.5), and 500 mM NaCl for 45 min at 37°C. For the RNase H control, cells were co-digested with both RNase A and RNase H. After digestion, cells were blocked with 5% BSA in PBS for 3 h at room temperature. Next, cells were incubated overnight at 4 °C with the anti-DNA-RNA hybrid (S9.6) antibody after washing with PBS, cells were incubated with a secondary antibody for 2 h. After washing with PBS, the coverslips were air-dried and mounted onto slides using 80% glycerol with 10 µg/mL Hoechst 33258.

### Detection of RNA/DNA hybrids through dot blot

Cells were seeded and treated with cisplatin and then lysed using DNAzol reagent and then genomic DNA was precipitated by adding 1 volume of 100% ethanol. The DNA pellet was washed once with 70% ethanol and resuspended in 1X NEB CutSmart buffer. To fragment the genomic DNA, a cocktail of restriction enzymes (EcoRI, HindIII, XbaI, and NotI) was added, and incubated overnight for complete digestion. Fragmented DNA was then extracted using a phenol-chloroform-isoamyl alcohol mixture (25:24:1). DNA was precipitated overnight at -20°C by adding 3M sodium acetate (1/10th volume) and 100% ethanol (2.5 volumes).

The DNA pellet was washed once with 70% ethanol and resuspended in TE buffer. DNA concentration was measured by nano-drop spectrophotometer, and 1 µg of DNA per sample were slot blotted onto nitrocellulose membranes and then probed with RNA-DNA hybrid (S9.6) antibody and then secondary antibody. For RNase H control sample, 1 µg sample was digested with RNase H for 6 h prior to loading. For loading control, DNA was probed with anti-DNA antibody (Sigma-Aldrich, St. Louis, MO).

### Chromatin bound RNA assay

Cells were seeded and then lysed using DNAzol reagent and then genomic DNA was precipitated by adding 1 volume of 100% ethanol. The precipitated genomic DNA was dissolved in TRIzol reagent to isolate the chromatin bound RNA and process the same for Real Time PCR. The equal amount of precipitated RNA (1 µg /sample) was used to prepare cDNA by using TOPscript™ cDNA Synthesis Kit. The specific primers were used to amplify the ATG7 at ORF region and at distal region of transcribing mRNA and then quantified by Real time PCR using SYBR green (from Biorad). The Ct values obtained for the desired amplicon in WRN^-/-^ were normalized with WRN^+/+.^

### Co-immunoprecipitation

For co-immunoprecipitation experiments, 5 × 10^6^ cells were seeded in 150 mm dishes. After 16 h, the cells were transfected with 15 µg of WRN-FLAG plasmid using the calcium chloride method. The cells were then incubated in fresh medium for 24 h before being harvested by scraping in PBS. The cell pellets were resuspended in IP lysis buffer (50 mM Tris-HCl, pH 8.0, 300 mM NaCl, 0.4% NP-40, protease inhibitor cocktail), supplemented with DNase I digestion buffer (1x final concentration from a 10x stock), 10 U/mL DNase I, and 5 U/mL benzonase. Lysis was performed on ice for 1 h with periodic vortexing. The lysate was centrifuge at 12,500 rpm for 20 min to remove cell debris, the supernatant was collected and resuspended in IP dilution buffer (50 mM Tris-HCl, pH 8.0, 0.4% NP-40, protease inhibitor cocktail). After determining the protein concentration by Bradford assay, 5% of lysate was saved as input sample and 1mg of protein per sample was incubated with anti-FLAG antibody at 4°C for overnight. After equilibrating the protein A/G magnetic beads, these beads were added in lysate and incubated for 4 h at 4°C with rotation. The beads were then washed, boiled in SDS sample buffer, and analyzed by Western blotting.

### RNA immunoprecipitation (RIP)

Ribonucleoprotein immunoprecipitation (RIP) was performed as described previously with minor modifications. Magnetic beads and antibody mix (anti WRN or IgG) was prepared before lysing the cells. U2-OS cells were harvested in ice old PBS using scrapper followed by lysis solution containing RNase inhibitor (20 units/ml). After centrifugation, supernatant was collected and subjected to DNase I digestion for 10 min. The lysate was then added to the antibody bounds beads and subjected to rotation for 4 h at 4°C. After 2 wash of the magnetic beads, 100 ml proteinase K was mixed into each IP sample. Acid phenol:chloroform was then used to separate out RNA from the samples, followed by addition of glycogen and ethanol to precipitate small amount of RNA. The RNA was then subjected to real time PCR using ATG7 primers.

### Patient cancer tissues qPCR

OriGene’s TissueScan qPCR Cancer Survey cDNA Array I (#CSRT101) was used to evaluate the gene expression of ATG7 (proximal and distal mRNA) and WRN across four different cancer types (lung, breast, kidney, and ovarian) by RT-qPCR using SYBR Green and normalized by normal sample, following the manufacturer’s protocol. For analysis, difference Ct values of proximal and distal mRNA was calculated and then normalized this and Ct value of WRN expression with normal patient sample Ct values.

### Statistical analysis

The data are presented as the mean ± S.E.M or mean ± S.D. Comparisons between two groups were performed using an unpaired Student’s t-test in GraphPad Prism 8.0 software. Comparisons between two groups were performed using one-way analysis of variance (ANOVA) and two-way ANOVA for multiple comparisons in GraphPad Prism 8.0 software. A value of *p*<0.05 was considered statistically significant.

## Primers used for the qPCR

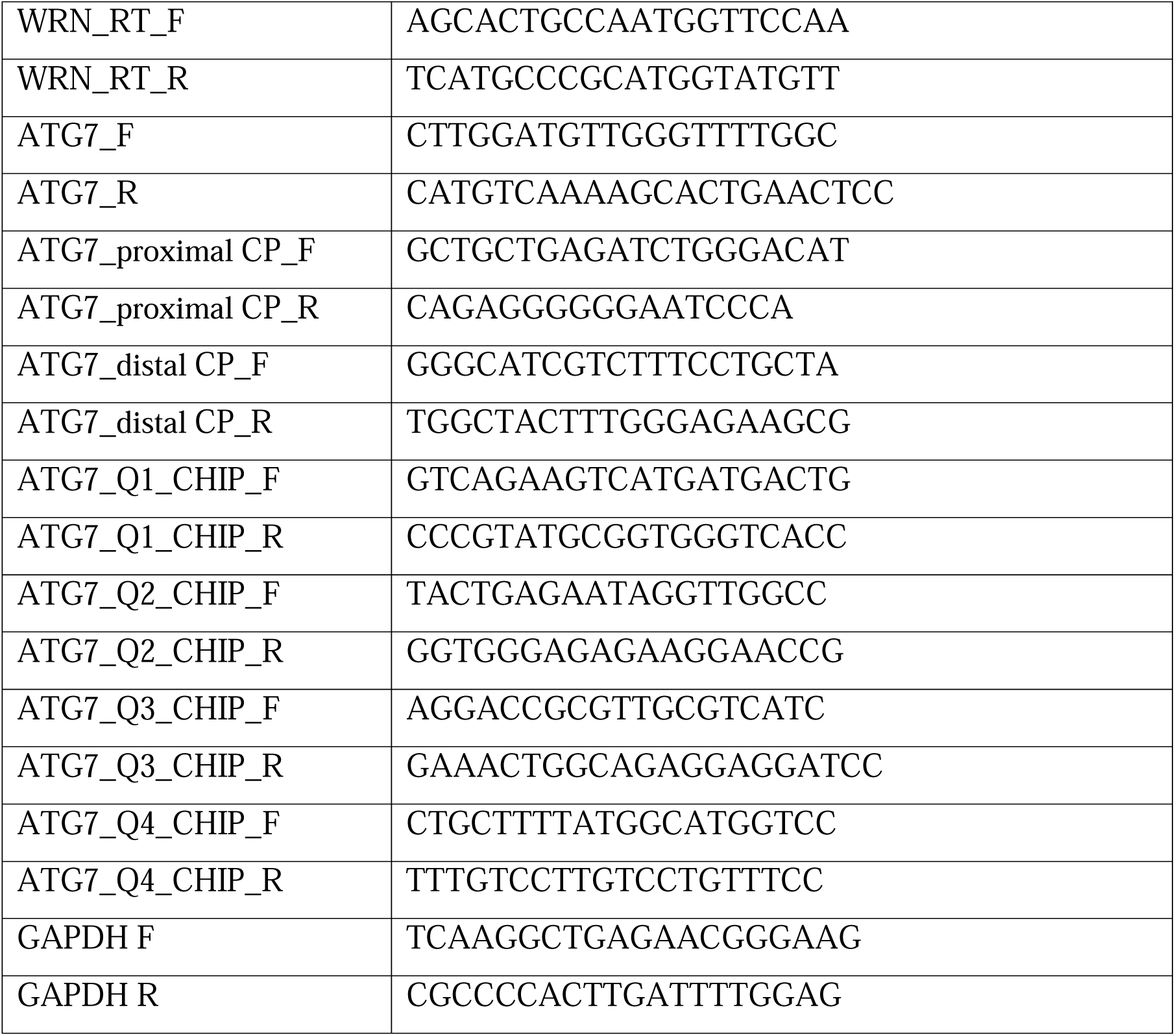

## Author contributions

Concept and design of the experiments: PG, MT (Tyagi), AGM, GPB and BSP; Execution of experiments: PG, MT (Tyagi), AGM, GPB, SC, SM, and BSP.; Experimental supports: MT (Megha), AC and SKB; Data analysis and curation: PG, MT (Tyagi), AGM, GPB, SKB and BSP; Manuscript draft writing: PG, MT (Tyagi) and BSP; Editing and finalizing the manuscript: All authors.

## References

(1) Muftuoglu, M.; Kusumoto, R.; Speina, E.; Beck, G.; Cheng, W.-H.; Bohr, V. A. Acetylation Regulates WRN Catalytic Activities and Affects Base Excision DNA Repair. PLOS ONE 2008, 3 (4), e1918. 10.1371/journal.pone.0001918.

(2) Agrelo, R.; Cheng, W.-H.; Setien, F.; Ropero, S.; Espada, J.; Fraga, M. F.; Herranz, M.; Paz, M. F.; Sanchez-Cespedes, M.; Artiga, M. J.; Guerrero, D.; Castells, A.; von Kobbe, C.; Bohr, V. A.; Esteller, M. Epigenetic Inactivation of the Premature Aging Werner Syndrome Gene in Human Cancer. Proc Natl Acad Sci U S A 2006, 103 (23), 8822–8827. 10.1073/pnas.0600645103.

(3) Lee, J. H.; Kim, S. S.; Kim, M. S.; Yoo, N. J.; Lee, S. H. WRN, the Werner Syndrome Gene, Exhibits Frameshift Mutations in Gastric and Colorectal Cancers. Pathol Oncol Res 2017, 23 (2), 451–452. 10.1007/s12253-016-0173-3.

(4) Zimmer K; Puccini A; Xiu J; Baca Y, Spizzo G, Lenz HJ, Battaglin F, Goldberg RM, Grothey A, Shields AF, Salem ME, Marshall JL, Korn WM, Wolf D, Kocher F, Seeber A. WRN-Mutated Colorectal Cancer Is Characterized by a Distinct Genetic Phenotype. Cancers (Basel) 2020, 12 (5), 1319. 10.3390/cancers12051319.

(5) Savva C; Sadiq M; Sheikh O; Karim S, Trivedi S, Green AR, Rakha EA, Madhusudan S, Arora A. Werner Syndrome Protein Expression in Breast Cancer. Clin Breast Cancer 2021, 21 (1), 57–73.e7. 10.1016/j.clbc.2020.07.013.

(6) Shamanna, R. A.; Lu, H.; Croteau, D. L.; Arora, A.; Agarwal, D.; Ball, G.; Aleskandarany, M. A.; Ellis, I. O.; Pommier, Y.; Madhusudan, S.; Bohr, V. A. Camptothecin Targets WRN Protein: Mechanism and Relevance in Clinical Breast Cancer. Oncotarget 2016, 7 (12), 13269–13284. 10.18632/oncotarget.7906.

(7) Gray M; Shen JC; Kamath-Loeb A; Blank A, Sopher BL, Martin GM, Oshima J, Loeb LA . The Werner Syndrome Protein Is a DNA Helicase. Nat Genet 1997, 17, 100–103. 10.1038/ng0997-100.

(8) Edwards D; Machwe A; Chen L; et al. The DNA Structure and Sequence Preferences of WRN Underlie Its Function in Telomeric Recombination Events. Nat Commun 2015, 6, 8331. 10.1038/ncomms9331.

(9) Rossi ML; Ghosh AK; Bohr VA. Roles of Werner Syndrome Protein in Protection of Genome Integrity. DNA Repair (Amst) 2010, 9 (3), 331–344. 10.1016/j.dnarep.2009.12.011.

(10) Datta A; Biswas K; Sommers JA; et al. WRN Helicase Safeguards Deprotected Replication Forks in BRCA2-Mutated Cancer Cells. Nat Commun 2021, 12, 6561. 10.1038/s41467-021-26811-w.

(11) Noto A; Valenzisi P; Di Feo F; et al. Phosphorylation-Dependent WRN-RPA Interaction Promotes Recovery of Stalled Forks at Secondary DNA Structure. Nat Commun 2025, 16, 997. 10.1038/s41467-025-55958-z.

(12) Patro BS; Frøhlich R; Bohr VA; Stevnsner T. WRN Helicase Regulates the ATR-CHK1-Induced S-Phase Checkpoint Pathway in Response to Topoisomerase-I-DNA Covalent Complexes. J Cell Sci 2011, 124 (Pt 23), 3967–3979. 10.1242/jcs.081372.

(13) Gupta, P.; Majumdar, A. G.; Patro, B. S. Non-Enzymatic Function of WRN RECQL Helicase Regulates Removal of Topoisomerase-I-DNA Covalent Complexes and Triggers NF-κB Signaling in Cancer. Aging Cell 2022, 21 (6), e13625. 10.1111/acel.13625.

(14) Chan EM; Shibue T; McFarland JM; Gaeta B, Ghandi M, Dumont N, Gonzalez A, McPartlan JS, Li T, Zhang Y, Bin Liu J, Lazaro JB, Gu P, Piett CG, Apffel A, Ali SO, Deasy R, Keskula P, Ng RWS, Roberts EA, Reznichenko E, Leung L, Alimova M, Schenone M, Islam M, Maruvka YE, Liu Y, Roper J, Raghavan S, Giannakis M, Tseng YY, Nagel ZD, D’Andrea A, Root DE, Boehm JS, Getz G, Chang S, Golub TR, Tsherniak A, Vazquez F, Bass AJ.. WRN Helicase Is a Synthetic Lethal Target in Microsatellite Unstable Cancers. Nature 2019, 568 (7753), 551–556. 10.1038/s41586-019-1102-x.

(15) Lieb, S.; Blaha-Ostermann, S.; Kamper, E.; Rippka, J.; Schwarz, C.; Ehrenhöfer-Wölfer, K.; Schlattl, A.; Wernitznig, A.; Lipp, J. J.; Nagasaka, K.; van der Lelij, P.; Bader, G.; Koi, M.; Goel, A.; Neumüller, R. A.; Peters, J.-M.; Kraut, N.; Pearson, M. A.; Petronczki, M.; Wöhrle, S. Werner Syndrome Helicase Is a Selective Vulnerability of Microsatellite Instability-High Tumor Cells. eLife 2019, 8, e43333. 10.7554/eLife.43333.

(16) Hao S; Tong J; Jha A; Risnik, D. Lizardo, X. Lu, A. Goel, P.L. Opresko, J. Yu, & L. Zhang. Synthetic Lethality of Werner Helicase and Mismatch Repair Deficiency Is Mediated by P53 and PUMA in Colon Cancer. Proc Natl Acad Sci U S A 2022, 119 (51), e2211775119. 10.1073/pnas.2211775119.

(17) Ferretti, S.; Hamon, J.; de Kanter, R.; Scheufler, C.; Andraos-Rey, R.; Barbe, S.; Bechter, E.; Blank, J.; Bordas, V.; Dammassa, E.; Decker, A.; Di Nanni, N.; Dourdoigne, M.; Gavioli, E.; Hattenberger, M.; Heuser, A.; Hemmerlin, C.; Hinrichs, J.; Kerr, G.; Laborde, L.; Jaco, I.; Núñez, E. J.; Martus, H.-J.; Quadt, C.; Reschke, M.; Romanet, V.; Schaeffer, F.; Schoepfer, J.; Schrapp, M.; Strang, R.; Voshol, H.; Wartmann, M.; Welly, S.; Zécri, F.; Hofmann, F.; Möbitz, H.; Cortés-Cros, M. Discovery of WRN Inhibitor HRO761 with Synthetic Lethality in MSI Cancers. Nature 2024, 629 (8011), 443–449. 10.1038/s41586-024-07350-y.

(18) Fang, E. F.; Hou, Y.; Lautrup, S.; Jensen, M. B.; Yang, B.; SenGupta, T.; Caponio, D.; Khezri, R.; Demarest, T. G.; Aman, Y.; Figueroa, D.; Morevati, M.; Lee, H.-J.; Kato, H.; Kassahun, H.; Lee, J.-H.; Filippelli, D.; Okur, M. N.; Mangerich, A.; Croteau, D. L.; Maezawa, Y.; Lyssiotis, C. A.; Tao, J.; Yokote, K.; Rusten, T. E.; Mattson, M. P.; Jasper, H.; Nilsen, H.; Bohr, V. A. NAD+ Augmentation Restores Mitophagy and Limits Accelerated Aging in Werner Syndrome. Nat Commun 2019, 10 (1), 5284. 10.1038/s41467-019-13172-8.

(19) Massip, L.; Garand, C.; Paquet, E. R.; Cogger, V. C.; O’Reilly, J. N.; Tworek, L.; Hatherell, A.; Taylor, C. G.; Thorin, E.; Zahradka, P.; Le Couteur, D. G.; Lebel, M. Vitamin C Restores Healthy Aging in a Mouse Model for Werner Syndrome. FASEB J 2010, 24 (1), 158–172. 10.1096/fj.09-137133.

(20) Maity, J.; Bohr, V. A.; Laskar, A.; Karmakar, P. Transient Overexpression of Werner Protein Rescues Starvation Induced Autophagy in Werner Syndrome Cells. Biochim Biophys Acta 2014, 1842 (12 Pt A), 2387–2394. 10.1016/j.bbadis.2014.09.007.

(21) Maity, J.; Das, B.; Bohr, V. A.; Karmakar, P. Acidic Domain of WRNp Is Critical for Autophagy and Up-Regulates Age Associated Proteins. DNA Repair (Amst) 2018, 68, 1–11. 10.1016/j.dnarep.2018.05.003.

(22) Li, B.; Iglesias-Pedraz, J. M.; Chen, L.-Y.; Yin, F.; Cadenas, E.; Reddy, S.; Comai, L. Downregulation of the Werner Syndrome Protein Induces a Metabolic Shift That Compromises Redox Homeostasis and Limits Proliferation of Cancer Cells. Aging Cell 2014, 13 (2), 367–378. 10.1111/acel.12181.

(23) Iglesias-Pedraz, J. M.; Fossatti-Jara, D. M.; Valle-Riestra-Felice, V.; Cruz-Visalaya, S. R.; Ayala Felix, J. A.; Comai, L. WRN Modulates Translation by Influencing Nuclear mRNA Export in HeLa Cancer Cells. BMC Mol Cell Biol 2020, 21 (1), 71. 10.1186/s12860-020-00315-9.

(24) Kadam, A.; Jubin, T.; Roychowdhury, R.; Begum, R. Role of PARP-1 in Mitochondrial Homeostasis. Biochim Biophys Acta Gen Subj 2020, 1864 (10), 129669. 10.1016/j.bbagen.2020.129669.

(25) Mishra, S. R.; Mishra, P.; Mahapatra, K. K.; Behera, B. P.; Kendre, G.; Alotaibi, M. R.; Pandey, V.; Patro, B. S.; Klionsky, D. J.; Bhutia, S. K. PPA2 Activates MTFP1-DNM1L Fission Signaling to Govern Mitochondrial Proliferation and Mitophagy. Autophagy 2025, 1–24. 10.1080/15548627.2025.2552900.

(26) Praharaj, P. P.; Patro, B. S.; Bhutia, S. K. Dysregulation of Mitophagy and Mitochondrial Homeostasis in Cancer Stem Cells: Novel Mechanism for Anti-Cancer Stem Cell-Targeted Cancer Therapy. Br J Pharmacol 2022, 179 (22), 5015–5035. 10.1111/bph.15401.

(27) Kanai, F.; Liu, H.; Field, S. J.; Akbary, H.; Matsuo, T.; Brown, G. E.; Cantley, L. C.; Yaffe, M. B. The PX Domains of P47phox and P40phox Bind to Lipid Products of PI(3)K. Nat Cell Biol 2001, 3 (7), 675–678. 10.1038/35083070.

(28) Rojansky, R.; Cha, M.-Y.; Chan, D. C. Elimination of Paternal Mitochondria in Mouse Embryos Occurs through Autophagic Degradation Dependent on PARKIN and MUL1. Elife 2016, 5, e17896. 10.7554/eLife.17896.

(29) Pai Bellare, G.; Saha, B.; Patro, B. S. Targeting Autophagy Reverses de Novo Resistance in Homologous Recombination Repair Proficient Breast Cancers to PARP Inhibition. Br J Cancer 2021, 124 (7), 1260–1274. 10.1038/s41416-020-01238-0.

(30) Onodera, J.; Ohsumi, Y. Autophagy Is Required for Maintenance of Amino Acid Levels and Protein Synthesis under Nitrogen Starvation. J Biol Chem 2005, 280 (36), 31582–31586. 10.1074/jbc.M506736200.

(31) Collier, J. J.; Suomi, F.; Oláhová, M.; McWilliams, T. G.; Taylor, R. W. Emerging Roles of ATG7 in Human Health and Disease. EMBO Mol Med 2021, 13 (12), e14824. 10.15252/emmm.202114824.

(32) Cogoi, S.; Xodo, L. E. G-Quadruplex Formation within the Promoter of the KRAS Proto-Oncogene and Its Effect on Transcription. Nucleic Acids Res 2006, 34 (9), 2536–2549. 10.1093/nar/gkl286.

(33) Siddiqui-Jain, A.; Grand, C. L.; Bearss, D. J.; Hurley, L. H. Direct Evidence for a G-Quadruplex in a Promoter Region and Its Targeting with a Small Molecule to Repress c-MYC Transcription. Proc Natl Acad Sci U S A 2002, 99 (18), 11593–11598. 10.1073/pnas.182256799.

(34) Ketkar, A.; Voehler, M.; Mukiza, T.; Eoff, R. L. Residues in the RecQ C-Terminal Domain of the Human Werner Syndrome Helicase Are Involved in Unwinding G-Quadruplex DNA. J Biol Chem 2017, 292 (8), 3154–3163. 10.1074/jbc.M116.767699.

(35) Johnson, J. E.; Cao, K.; Ryvkin, P.; Wang, L.-S.; Johnson, F. B. Altered Gene Expression in the Werner and Bloom Syndromes Is Associated with Sequences Having G-Quadruplex Forming Potential. Nucleic Acids Res 2010, 38 (4), 1114–1122. 10.1093/nar/gkp1103.

(36) Moruno-Manchon, J. F.; Koellhoffer, E. C.; Gopakumar, J.; Hambarde, S.; Kim, N.; McCullough, L. D.; Tsvetkov, A. S. The G-Quadruplex DNA Stabilizing Drug Pyridostatin Promotes DNA Damage and Downregulates Transcription of Brca1 in Neurons. Aging (Albany NY) 2017, 9 (9), 1957–1970. 10.18632/aging.101282.

(37) Kikin, O.; D’Antonio, L.; Bagga, P. S. QGRS Mapper: A Web-Based Server for Predicting G-Quadruplexes in Nucleotide Sequences. Nucleic Acids Res 2006, 34 (Web Server issue), W676–682. 10.1093/nar/gkl253.

(38) Park, S. M.; Ou, J.; Chamberlain, L.; Simone, T. M.; Yang, H.; Virbasius, C.-M.; Ali, A. M.; Zhu, L. J.; Mukherjee, S.; Raza, A.; Green, M. R. U2AF35(S34F) Promotes Transformation by Directing Aberrant ATG7 Pre-mRNA 3’ End Formation. Mol Cell 2016, 62 (4), 479–490. 10.1016/j.molcel.2016.04.011.

(39) Millevoi, S.; Loulergue, C.; Dettwiler, S.; Karaa, S. Z.; Keller, W.; Antoniou, M.; Vagner, S. An Interaction between U2AF 65 and CF I(m) Links the Splicing and 3’ End Processing Machineries. EMBO J 2006, 25 (20), 4854–4864. 10.1038/sj.emboj.7601331.

(40) Guo, J. Y.; Karsli-Uzunbas, G.; Mathew, R.; Aisner, S. C.; Kamphorst, J. J.; Strohecker, A. M.; Chen, G.; Price, S.; Lu, W.; Teng, X.; Snyder, E.; Santanam, U.; Dipaola, R. S.; Jacks, T.; Rabinowitz, J. D.; White, E. Autophagy Suppresses Progression of K-Ras-Induced Lung Tumors to Oncocytomas and Maintains Lipid Homeostasis. Genes Dev 2013, 27 (13), 1447–1461. 10.1101/gad.219642.113.

(41) Tang, W.; Robles, A. I.; Beyer, R. P.; Gray, L. T.; Nguyen, G. H.; Oshima, J.; Maizels, N.; Harris, C. C.; Monnat, R. J. The Werner Syndrome RECQ Helicase Targets G4 DNA in Human Cells to Modulate Transcription. Hum Mol Genet 2016, 25 (10), 2060–2069. 10.1093/hmg/ddw079.

(42) Marabitti, V.; Lillo, G.; Malacaria, E.; Palermo, V.; Sanchez, M.; Pichierri, P.; Franchitto, A. ATM Pathway Activation Limits R-Loop-Associated Genomic Instability in Werner Syndrome Cells. Nucleic Acids Res 2019, 47 (7), 3485–3502. 10.1093/nar/gkz025.

(43) Chen, S.; Wang, Z.; Cao, Z.; Xu, M.; Zhang, Y. Targeting the Werner Syndrome Protein in Microsatellite Instability Cancers: Mechanisms and Therapeutic Potential. Clin Exp Med 2025, 25 (1), 278. 10.1007/s10238-025-01781-1.

(44) Behan, F. M.; Iorio, F.; Picco, G.; Gonçalves, E.; Beaver, C. M.; Migliardi, G.; Santos, R.; Rao, Y.; Sassi, F.; Pinnelli, M.; Ansari, R.; Harper, S.; Jackson, D. A.; McRae, R.; Pooley, R.; Wilkinson, P.; van der Meer, D.; Dow, D.; Buser-Doepner, C.; Bertotti, A.; Trusolino, L.; Stronach, E. A.; Saez-Rodriguez, J.; Yusa, K.; Garnett, M. J. Prioritization of Cancer Therapeutic Targets Using CRISPR-Cas9 Screens. Nature 2019, 568 (7753), 511–516. 10.1038/s41586-019-1103-9.

(45) Kategaya, L.; Perumal, S. K.; Hager, J. H.; Belmont, L. D. Werner Syndrome Helicase Is Required for the Survival of Cancer Cells with Microsatellite Instability. iScience 2019, 13, 488–497. 10.1016/j.isci.2019.02.006.

(46) McDonald ER 3rd; de Weck A; Schlabach MR; et al. Project DRIVE: A Compendium of Cancer Dependencies and Synthetic Lethal Relationships Uncovered by Large-Scale, Deep RNAi Screening. Cell 2017, 170 (3), 577–592.e10. 10.1016/j.cell.2017.07.005.

(47) Cowley, G. S.; Weir, B. A.; Vazquez, F.; Tamayo, P.; Scott, J. A.; Rusin, S.; East-Seletsky, A.; Ali, L. D.; Gerath, W. F.; Pantel, S. E.; Lizotte, P. H.; Jiang, G.; Hsiao, J.; Tsherniak, A.; Dwinell, E.; Aoyama, S.; Okamoto, M.; Harrington, W.; Gelfand, E.; Green, T. M.; Tomko, M. J.; Gopal, S.; Wong, T. C.; Li, H.; Howell, S.; Stransky, N.; Liefeld, T.; Jang, D.; Bistline, J.; Hill Meyers, B.; Armstrong, S. A.; Anderson, K. C.; Stegmaier, K.; Reich, M.; Pellman, D.; Boehm, J. S.; Mesirov, J. P.; Golub, T. R.; Root, D. E.; Hahn, W. C. Parallel Genome-Scale Loss of Function Screens in 216 Cancer Cell Lines for the Identification of Context-Specific Genetic Dependencies. Scientific Data 2014, 1 (1), 140035. 10.1038/sdata.2014.35.

(48) Baltgalvis, K. A.; Lamb, K. N.; Symons, K. T.; Wu, C.-C.; Hoffman, M. A.; Snead, A. N.; Song, X.; Glaza, T.; Kikuchi, S.; Green, J. C.; Rogness, D. C.; Lam, B.; Rodriguez-Aguirre, M. E.; Woody, D. R.; Eissler, C. L.; Rodiles, S.; Negron, S. M.; Bernard, S. M.; Tran, E.; Pollock, J.; Tabatabaei, A.; Contreras, V.; Williams, H. N.; Pastuszka, M. K.; Sigler, J. J.; Pettazzoni, P.; Rudolph, M. G.; Classen, M.; Brugger, D.; Claiborne, C.; Plancher, J.-M.; Cuartas, I.; Seoane, J.; Burgess, L. E.; Abraham, R. T.; Weinstein, D. S.; Simon, G. M.; Patricelli, M. P.; Kinsella, T. M. Chemoproteomic Discovery of a Covalent Allosteric Inhibitor of WRN Helicase. Nature 2024, 629 (8011), 435–442. 10.1038/s41586-024-07318-y.

(49) Robinson, J.; Raguseo, F.; Nuccio, S. P.; Liano, D.; Di Antonio, M. DNA G-Quadruplex Structures: More than Simple Roadblocks to Transcription? Nucleic Acids Res 2021, 49 (15), 8419–8431. 10.1093/nar/gkab609.

(50) Hänsel-Hertsch, R.; Spiegel, J.; Marsico, G.; Tannahill, D.; Balasubramanian, S. Genome-Wide Mapping of Endogenous G-Quadruplex DNA Structures by Chromatin Immunoprecipitation and High-Throughput Sequencing. Nat Protoc 2018, 13 (3), 551–564. 10.1038/nprot.2017.150.

(51) Mendoza, O.; Bourdoncle, A.; Boulé, J.-B.; Brosh, R. M.; Mergny, J.-L. G-Quadruplexes and Helicases. Nucleic Acids Res 2016, 44 (5), 1989–2006. 10.1093/nar/gkw079.

(52) Palangat, M.; Anastasakis, D. G.; Fei, D. L.; Lindblad, K. E.; Bradley, R.; Hourigan, C. S.; Hafner, M.; Larson, D. R. The Splicing Factor U2AF1 Contributes to Cancer Progression through a Noncanonical Role in Translation Regulation. Genes Dev 2019, 33 (9–10), 482–497. 10.1101/gad.319590.118.

(53) Feng, Y.; Zhang, H.; Han, J.; Cui, B.; Qin, L.; Zhang, L.; Li, Q.; Wu, X.; Xiao, N.; Zhang, Y.; Lin, T.; Liu, H.; Sun, T. HSF4/COIL Complex-Dependent R-Loop Mediates Ultraviolet-Induced Inflammatory Skin Injury. Clinical and Translational Medicine 2023, 13 (7), e1336. 10.1002/ctm2.1336.

(54) Long, J. S.; Kania, E.; McEwan, D. G.; Barthet, V. J. A.; Brucoli, M.; Ladds, M. J. G. W.; Nössing, C.; Ryan, K. M. ATG7 Is a Haploinsufficient Repressor of Tumor Progression and Promoter of Metastasis. Proceedings of the National Academy of Sciences 2022, 119 (28), e2113465119. 10.1073/pnas.2113465119.

(55) Gupta, P.; Saha, B.; Chattopadhyay, S.; Patro, B. S. Pharmacological Targeting of Differential DNA Repair, Radio-Sensitizes WRN-Deficient Cancer Cells in Vitro and in Vivo. Biochem Pharmacol 2021, 186, 114450. 10.1016/j.bcp.2021.114450.

(56) Hänsel-Hertsch, R.; Spiegel, J.; Marsico, G.; Tannahill, D.; Balasubramanian, S. Genome-Wide Mapping of Endogenous G-Quadruplex DNA Structures by Chromatin Immunoprecipitation and High-Throughput Sequencing. Nature Protocols 2018, 13 (3), 551–564. 10.1038/nprot.2017.150.

