## Supplementary Figures for "WRN helicase upregulates mitophagy by resolving an intricate nexus of G-quadruplexes-R loops-ATG7 pre-mRNA maturation in cancer"

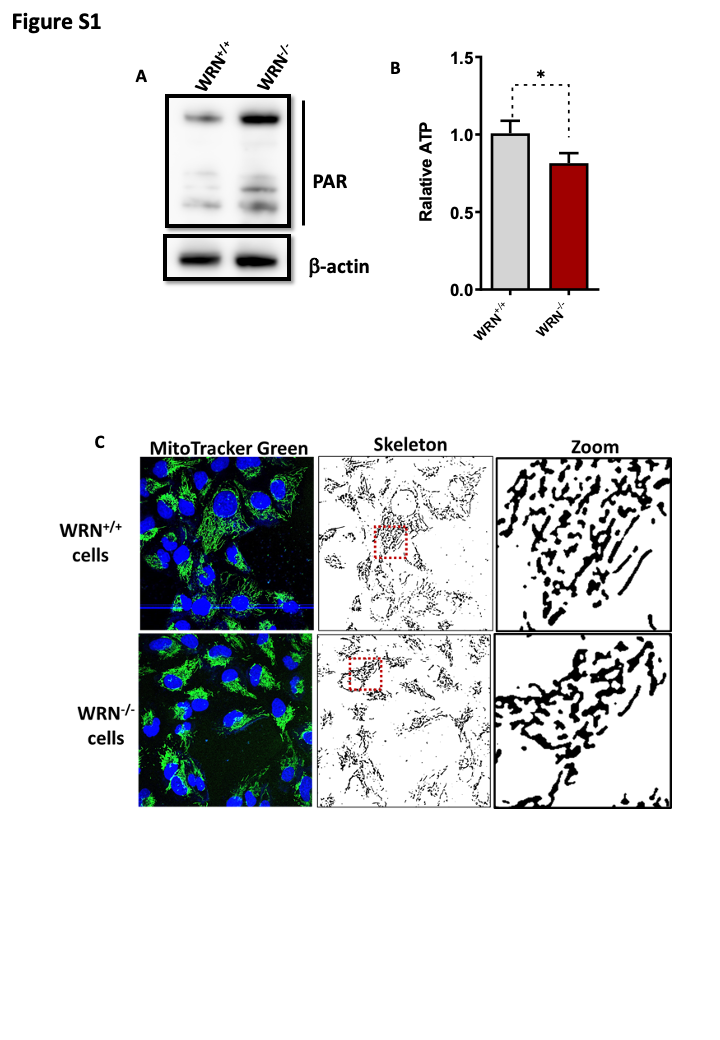

**Figure S1. PAR, ATP level and mitochondrial morphology (skeleton) in WRN^+/+^ and WRN^-/-^.** (A) PAR levels were assessed by Western blotting in WRN^+/+^ and WRN^-/-^ cells, probing for PAR and using β-actin as a loading control. (B) Relative ATP levels were measured in WRN^+/+^ and WRN^-/-^ cells using an ATP assay kit. (C) Representative confocal images of WRN^+/+^ and WRN^-/-^ cells stained with MitoTracker Green for mitochondrial morphology analysis. Zoomed-in images show the mitochondrial structure in both cell types. Skeleton of mitochondria and analysis is performed by Fiji software. All the values indicated are mean ± SEM (n = 3), **p* < 0.05.

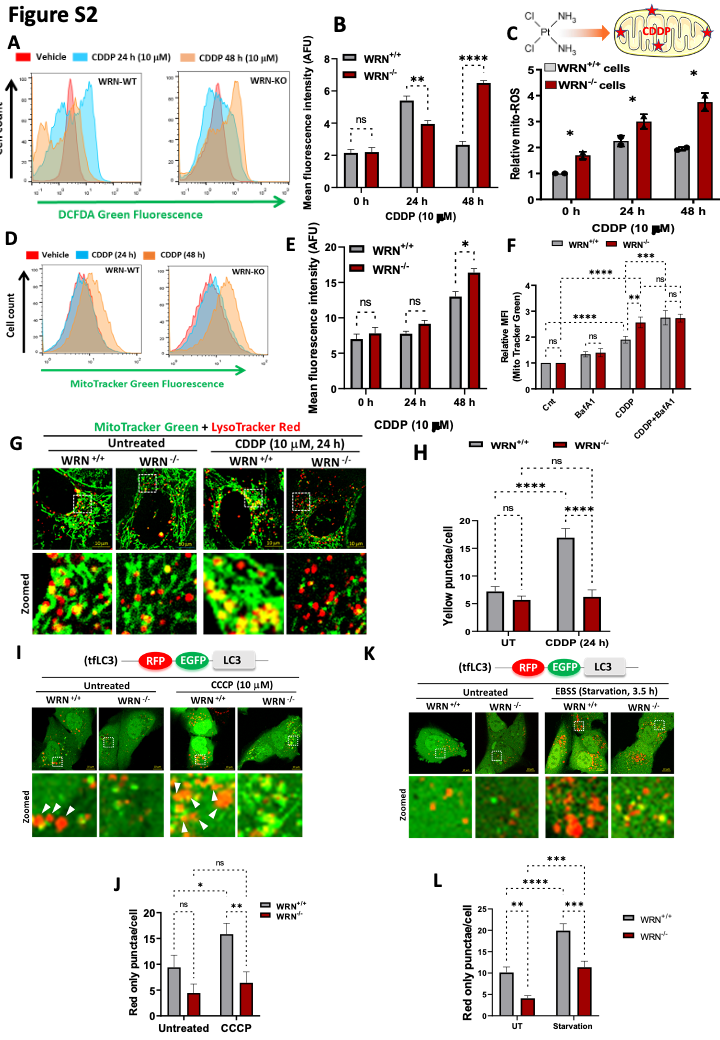

**Figure S2. WRN affects mitochondrial and autophagic responses to cisplatin (CDDP) and starvation.** (A, B) Flow cytometric analysis of DCFDA green fluorescence, for detaction of reactive oxygen species (ROS), in WRN^+/+^ and WRN^-/-^ cells treated with vehicle or CDDP. Mean DCFDA fluorescence intensity (AU) was quantified. (C) Relative mitochondrial ROS (mito-ROS) levels in WRN^+/+^ and WRN-/- cells, in response to CDDP treatment, was assessed by flow cytometry. (D, E) Flow cytometric analysis was carried out to assess MitoTracker Green fluorescence in WRN^+/+^ and WRN^-/-^ cells in untreated and CDDP-treated cells. (F) WRN^+/+^ and WRN^-/-^ cells were untreated or treated with CDDP (10 μM, 24 h) in the absence or presence of bafilomycin (BafA1). Cells were treated with BafA1 during last 3 h of CDDP treatment. Cells were stained with MitoTracker Green and relative mean fluorescence intensity (MFI) of MitoTracker Green was assessed using flow cytometry. (G, H) Cells were untreated or treated with CDDP (10 μM, 24 h) and stained mitochondria and lysosomes were stained MitoTracker Green and LysoTracker Red, respectively. Confocal images were acquired to analyze the co-localization of mitochondria and lysosomes (yellow puncta) in WRN^+/+^ and WRN^-/-^ cells. Zoomed-in images highlight the co-localization of mitochondria and lysosomes. Yellow puncta (representing co-localization of mitochondria and lysosomes) per WRN^+/+^ and WRN^-/-^ cell was quantified. (I, J) Cells, expressing tf-LC3, were untreated or treated with CDDP (10 μM, 24 h). Confocal images were acquired to assess autophagic flux. Red only punctae, representing autophagic flux, per cell was quantified. Zoomed-in images show the level of autophagic flux in WRN^+/+^ and WRN^-/-^ cells. (K, L) Cells, expressing tf-LC3, were untreated or starved in EBSS medium (3.5 h). Confocal images were acquired to assess autophagic flux. Red only punctae, representing autophagic flux, per cell was quantified. Zoomed-in images show the level of autophagic flux in WRN^+/+^ and WRN^-/-^ cells. All the values indicated are mean ± SEM (n = 3-4), ns denotes non-significant, **p* < 0.05, ***p* < 0.001, ****p* < 0.001 and *****p* < 0.0001.

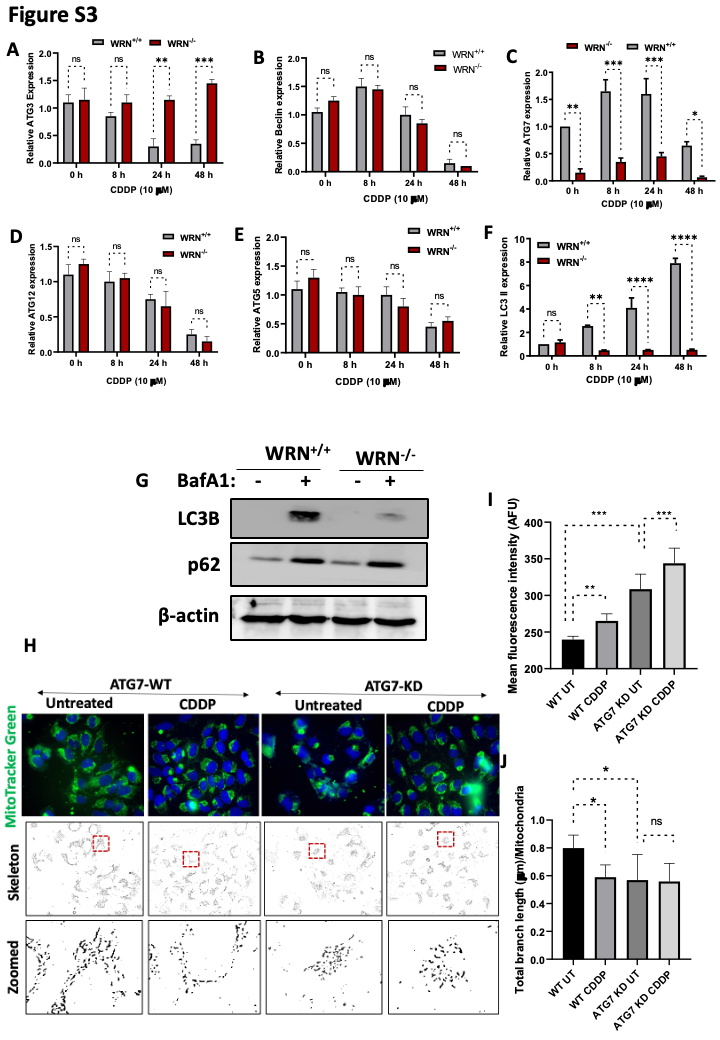

**Figure S3. Role of WRN and ATG7 in autophagy and mitochondrial dynamics.** (A-F) Relative expression of autophagosome regulating proteins in WRN^+/+^ and WRN^-/-^ cells in response to CDDP treatment was assessed by western blotting. (G) Cells were untreated or treated with BafA1 for 3 h. LC3-II and p62 were assessed using western blotting. (H-J) WT and ATG7-KD U2-OS cells were treated with CDDP (10 μM, 24 h). Cells were stained with MitoTracker Green and confocal images were acquired. Total mitochondrial mass was analysed by quantifying mean fluorescence intensity of MitoTracker Green. Mitochondrial morphology and branching were assessed staining followed by skeletonizing of mitochondrial structures (mitoTracker Green) in WRN^+/+^ and WRN^-/-^ cells. Representative images are shown. Total branch length (µm) of mitochondria in ATG7-WT and ATG7-KD cells was quantified. All the values indicated are mean ± SEM (n = 3-4), ns denotes non-significant, **p* < 0.05, ***p* < 0.001, ****p* < 0.001 and *****p* < 0.0001.
